# Crosstalk between chloroplast protein import and the SUMO system revealed through genetic and molecular investigation

**DOI:** 10.1101/2020.07.21.213355

**Authors:** Samuel James Watson, Na Li, Yiting Ye, Feijie Wu, Qihua Ling, R. Paul Jarvis

## Abstract

The chloroplast proteome contains thousands of different proteins that are encoded by the nuclear genome. These proteins are imported into the chloroplast via the action of the TOC translocase and associated downstream systems. Our recent work has revealed that the stability of the TOC complex is dynamically regulated by the ubiquitin-dependent chloroplast-associated protein degradation (CHLORAD) pathway. Here, we demonstrate that the stability of the TOC complex is also regulated by the SUMO system. *Arabidopsis* mutants representing almost the entire SUMO conjugation pathway can partially suppress the phenotype of *ppi1*, a pale yellow mutant lacking the Toc33 protein. This suppression is linked to the increased stability of TOC proteins and improvements in chloroplast development. In addition, we demonstrate using molecular and biochemical experiments that the SUMO system directly targets TOC proteins. Thus, we have identified a regulatory link between the SUMO system and chloroplast protein import.

## Introduction

The chloroplast is a membrane-bound organelle that houses photosynthesis in all green plants (Jarvis and Lopez-Juez, 2013). Chloroplasts have an unusual evolutionary history – they are the integrated descendants of a cyanobacterial ancestor that entered the eukaryotic lineage via endosymbiosis. Although chloroplasts retain small genomes, almost all of the proteins required for chloroplast development and function are now encoded by the central, nuclear genome (Jarvis, 2008). These proteins must be imported into the organelle after synthesis in the cytosol, and this import is mediated by the coordinate action of the TOC and TIC complexes (the translocons at the outer and inner envelope membranes of chloroplasts) (Jarvis, 2008).

The TOC complex contains three major components: the Omp85 (outer membrane protein, 85 kDa)-related protein, Toc75, which serves as a membrane channel (Schnell et al., 1994; Tranel et al., 1995), and two GTPase-domain receptor proteins, Toc33 and Toc159 (Hirsch et al., 1994; Kessler et al., 1994; Perry and Keegstra, 1994; Jarvis et al., 1998; Jarvis, 2008). Toc33 and Toc159 project into the cytosol and bind incoming preproteins.

The key components of the TOC complex were identified more than two decades ago (Hirsch et al., 1994; Kessler et al., 1994; Schnell et al., 1994; Tranel et al., 1995; Jarvis, 2008). However, the regulation of the activity and stability of the complex was, until recently, poorly understood. Major insights came from a forward genetic screen for suppressors of the pale yellow Toc33 mutant, *ppi1* (Ling et al., 2012). In this screen, SP1 (SUPPRESSOR OF PPI1 LOCUS 1), a novel RING-type E3 ubiquitin ligase, was identified. A series of *sp1* mutations were shown to partially suppress the phenotypic defects of *ppi1* with respect to chlorosis, chloroplast development, and chloroplast protein import. In addition, SP1 function was shown to promote plastid interconversion events (for example, the development of the chloroplast from its precursor organelle, the etioplast). Later work demonstrated that SP1 function is also important for abiotic stress tolerance, by enabling optimisation of the organellar proteome via protein import regulation (Ling and Jarvis, 2015). Thus, through SP1, the ubiquitin-proteasome system promotes TOC complex degradation and reconfiguration in response to developmental and/or environmental stimuli.

Ubiquitinated TOC proteins are extracted from the chloroplast outer envelope membrane and degraded in the cytosol. Recent work identified two proteins that physically associate with SP1 and promote the membrane extraction of TOC proteins (Ling et al., 2019). These are SP2, an Omp85-type β-barrel channel protein that was identified in the same genetic screen as SP1, and Cdc48, a well-characterised cytosolic AAA+ chaperone ATPase that provides the motive force for the extraction of proteins from the chloroplast outer envelope. The three proteins – SP1, SP2 and Cdc48 – together define a new pathway for the ubiquitination, membrane extraction, and degradation of chloroplast outer envelope proteins, which has been named chloroplast-associated protein degradation, or CHLORAD. In addition to CHLORAD, there exist cytosolic ubiquitin-dependent systems that also contribute to chloroplast biogenesis, by regulating the levels of unimported preproteins (Lee et al., 2009; Grimmer et al., 2020), and by controlling the stability of the Toc159 receptor prior to its integration into the outer envelope membrane (Shanmugabalaji et al., 2018).

The discovery of SP1 and the CHLORAD pathway demonstrated that the TOC complex is not static but, instead, can be rapidly ubiquitinated and degraded in response to developmental and environmental stimuli. To complement this work, we decided to explore whether the TOC complex is also regulated by the SUMO system. This work was motivated by the results of a high-throughput screen for SUMO substrates in *Arabidopsis* (Elrouby and Coupland, 2010). This screen suggested that Toc159, a key component of the TOC complex, is a SUMO substrate. SUMOylation is intricately involved in plant development and stress adaptation, and so we were interested to determine whether the TOC complex is targeted by the SUMO system, and whether any such SUMOylation is functionally important. As crosstalk between the SUMO system and the ubiquitin-proteasome system is common, we reasoned that answering these questions might provide insights into the regulation of SP1 and the CHLORAD pathway.

To explore the relationship between chloroplast protein import and the SUMO system, we carried out a comprehensive series of genetic, molecular and biochemical experiments. Mutants representing most components of the *Arabidopsis* SUMO pathway were found to partially suppress the phenotype of the chlorotic Toc33 null mutant, *ppi1*, with respect to leaf chlorophyll accumulation, chloroplast development, and TOC protein abundance. Conversely, overexpression of either *SUMO1* or *SUMO3* enhanced the severity of the *ppi1* phenotype. Moreover, the E2 SUMO conjugating enzyme, SCE1, was found to physically interact with the TOC complex in bimolecular fluorescence complementation experiments; and TOC proteins were seen to physically associate with SUMO proteins in immunoprecipitation assays. In combination, our data conclusively demonstrate significant crosstalk between the SUMO system and chloroplast protein import, and emphasise the complexity of the regulation of the TOC translocase.

## Results

### The E2 SUMO conjugating enzyme mutant *sce1-4*, and the E3 SUMO ligase mutants *siz1-4* and *siz1-5*, partially suppress the phenotype of Toc33 mutant, *ppi1*

Two key components of the CHLORAD pathway, SP1 and SP2, were identified in a forward genetic screen for suppressors of *ppi1*, an *Arabidopsis* Toc33 null mutant (Ling et al., 2012; Ling et al., 2019). Both *sp1* and *sp2* mutants can partially suppress the *ppi1* phenotype with respect to chlorophyll accumulation, chloroplast development, and TOC protein abundance. To investigate whether the TOC complex is targeted by the SUMO system, we obtained several *Arabidopsis* SUMO system mutants, crossed them with *ppi1*, and carefully examined the phenotypes of the resulting double mutants. This reverse genetic approach was possible because the basic architecture of the *Arabidopsis* SUMO system is remarkably simple. In the SUMO pathway, the ubiquitin-like SUMO modifier protein is conjugated to substates by the coordinated action of E1 activating enzymes, E2 conjugating enzymes, and E3 SUMO ligases. Although thousands of proteins are SUMOylated in *Arabidopsis*, there is just one known E1 SUMO activating enzyme, one known E2 SUMO conjugating enzyme, and only two known E3 SUMO ligases of canonical function (Saracco et al., 2007; Ishida et al., 2009).

First, we analysed *sce1-4*, a weak mutant allele of the sole E2 SUMO conjugating enzyme gene in *Arabidopsis*, which is an essential gene (Saracco et al., 2007). The *sce1-4* mutant shows a moderate reduction in the expression of SCE1 and in global levels of SUMOylation, but it displays no obvious visible phenotypic defects under steady-state conditions (Saracco et al., 2007). The *ppi1 sce1-4* double mutant was phenotypically characterised, and, intriguingly, it appeared greener than the *ppi1* single mutant (Figure 1A; Figure 1 – Supplement 1A). This was linked to a moderate increase in leaf chlorophyll concentration (Figure 1B; Figure 1 – Supplement 1B). Next, we asked whether the phenotypic suppression observed in *ppi1 sce1-4* was linked to changes in the development of chloroplasts. The chloroplasts of *ppi1 sce1-4* were visualised via transmission electron microscopy. Interestingly, the chloroplasts of the *ppi1 sce1-4* double mutant appeared larger and better developed than those of the *ppi1* control (Figure 1C). The transmission electron micrographs were quantitatively analysed, and the *ppi1 sce1-4* chloroplasts were indeed found to be significantly larger than those of *ppi1* (Figure 1D), with larger, more interconnected thylakoidal granal stacks (Figures 1E and 1F).

**Figure 1.**
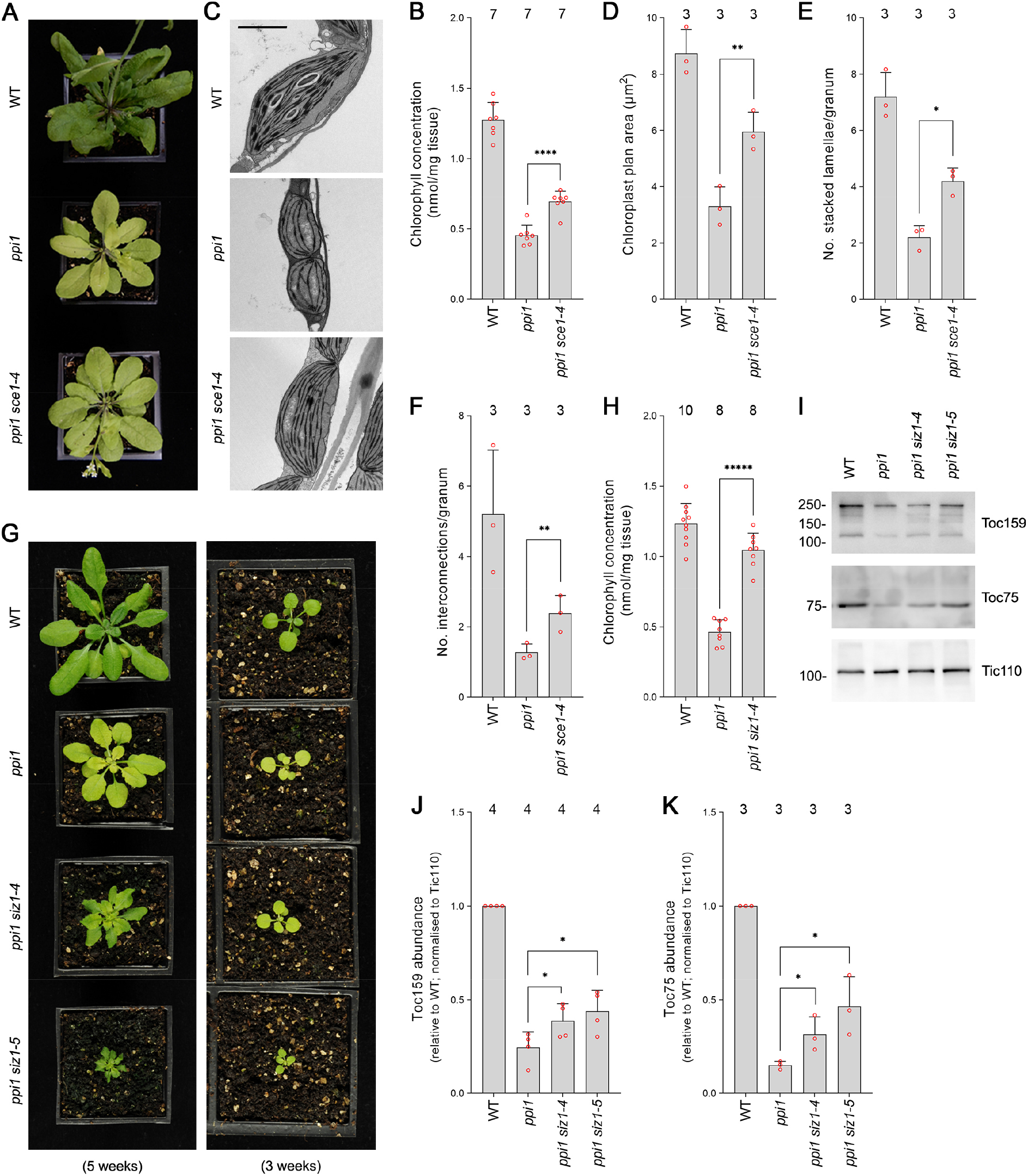
The E2 SUMO conjugating enzyme mutation, *sce1-4*, and the E3 SUMO ligase mutations, *siz1-4* and *siz1-5*, suppress the phenotype of plastid protein import mutant, *ppi1*. **(A)** The *ppi1 sce1-4* double mutant appeared greener than *ppi1* after approximately five weeks of growth on soil. **(B)** The *ppi1 sce1-4* double mutant showed enhanced accumulation of chlorophyll relative to *ppi1* after approximately five weeks of growth on soil. Measurements were taken from the plants shown in (A) on the day of photography, as well as additional similar plants. There were significant differences between the *ppi1* and *ppi1 sce1-4* plants (Two-tailed t-test, unpaired samples, T = 6.15, *p* = 0.000049). **(C)** Transmission electron microscopy revealed improved chloroplast development in mature rosette leaf mesophyll tissue of *ppi1 sce1-4* plants relative to *ppi1*. Plants that had been grown on soil for approximately four weeks were analysed, and representative images are shown. Scale bar = 2 μm. **(D)** Chloroplast plan area was elevated in *ppi1 sce1-4* relative to *ppi1*. The transmission electron microscopy dataset was quantified. There were significant differences between the *ppi1* and *ppi1 sce1-4* plants (Two-tailed t-test, unpaired samples, T = 4.65, *p* = 0.009674). **(E, F)** Thylakoid membrane development was increased in *ppi1 sce1-4* relative to *ppi1*. The number of stacked thylakoidal lamellae per granum (E), and the number of stromal thylakoidal lamellae emanating from each granum (granal interconnections) (F), was analysed using the transmission electron microscopy dataset. There were significant differences between the *ppi1* and *ppi1 sce1-4* plants (Two-tailed t-test, unpaired samples: T = 5.53, *p* = 0.005221 [E]; T = 3.38, *p* = 0.0277 [F]). **(G)** The *ppi1 siz1-4* and *ppi1 siz1-5* double mutants appeared greener than *ppi1* after different periods of growth on soil. The plants were photographed after three weeks of growth (right panel) and then again after five weeks of growth (left panel). **(H)** The *ppi1 siz1-4* double mutant showed enhanced accumulation of chlorophyll relative to *ppi1* after approximately five weeks of growth on soil. Measurements were taken from the plants shown in (G) on the day of photography, as well as additional similar plants. There were significant differences between the *ppi1* and *ppi1 siz1-4* plants (Two-tailed t-test, unpaired samples, T = 11.01, *p* < 0.00001). **(I)** TOC protein accumulation was improved in *ppi1 siz1-4* and *ppi1 siz1-5* relative to *ppi1*. Analysis of the levels of Toc75 and Toc159 in *ppi1 siz1-4*, *ppi1 siz1-5*, and relevant control plants was conducted by immunoblotting. Protein samples were taken from whole seedlings that had been grown on soil for approximately two weeks (the plants shown in Figure 1 – Supplement 1C on the day of photography). Tic110, a TIC-associated protein, was included as a compartment-specific loading control. Migration positions of standards are displayed to the left of the gel images, and sizes are indicated in kDa. Unprocessed membrane images are displayed in Source data 1. **(J, K)** Toc159 and Toc75 protein accumulation was improved in *ppi1 siz1-4* and *ppi1 siz1-5* relative to *ppi1*. Specific bands in (I) and in Source data 1 were quantified. Error bars indicate standard deviation from the mean. The numbers above the graph indicate the number of biological replicates per sample. There were significant differences between the *ppi1* and *ppi1 siz1-4* samples (One-tailed t-test, unpaired samples: T = 2.26, *p* = 0.032316 [J, Toc159]; T = 2.93, *p* = 0.021334 [K, Toc75]) and between the *ppi1* and *ppi1 siz1-5* samples (One-tailed t-test, unpaired samples: T = 2.76, *p* = 0.01639 [J, Toc159]; T = 3.36, *p* = 0.014118 [K, Toc75]). In all bar charts, error bars indicate standard deviation from the mean, and open red circles indicate individual data points. The numbers above the graphs indicate the number of biological replicates per sample. Statistical significance is indicated as follows: *, *p*<0.05; **, *p*<0.01; ****, *p*<0.0001; *****, *p*<0.00001.

SUMO conjugation is usually dependent on the action of E3 SUMO ligases. In *Arabidopsis*, the best characterised E3 SUMO ligase is SIZ1 (Kurepa et al., 2003; Miura et al., 2005; Saracco et al., 2007). SIZ1 is not essential, but null mutants display severely dwarfed phenotypes. In order to include SIZ1 in our genetic analysis, we obtained two new T-DNA insertion alleles, and named them *siz1-4* and *siz1-5.* While both mutants were visibly similar to the published mutants (Miura et al., 2005; Liu et al., 2019), *siz1-4* showed a milder phenotype with only moderate growth retardation when grown to maturity. We mapped the integration sites of the T-DNA insertions in these two mutants (Figure 1 – Supplement 2A), and showed that both display a strong reduction in *SIZ1* transcript by RT-PCR analysis (Figure 1 – Supplement 2B). In addition, both mutants displayed defects in global SUMOylation in response to heat shock, similar to the published alleles (Figure 1 – Supplement 3). The two new *siz1* mutants were crossed with *ppi1* and the resulting double mutants were phenotypically characterised. Both the *ppi1 siz1-4* and the *ppi1 siz1-5* double mutants appeared greener than the *ppi1* control (Figure 1G; Figure 1 – Supplement 1C). In addition, the double mutants showed dramatic increases in leaf chlorophyll concentration relative to *ppi1* (Figure 1H; Figure 1 – Supplement 1D). Next, we asked whether the phenotypic suppression observed in *ppi1 siz1-4* and *ppi1 siz1-5* was linked to changes in the abundance of TOC proteins. To this end, protein samples were taken from the two double mutants and relevant control plants and resolved via immunoblotting. Both double mutants displayed clear increases in the abundance of Toc159 and Toc75, two core components of the TOC complex, relative to *ppi1* (Figures 1I, 1J, and 1K).

### The suppression effects mediated by the SUMO system mutants are specific

As discussed in the previous section, the SUMO system is encoded by a remarkably small number of genes in *Arabidopsis*. As a consequence, SUMO system mutants have highly pleiotropic molecular and physiological phenotypes. We therefore asked whether the partial suppression of *ppi1* by SUMO system mutants was specific to the *ppi1* background. We crossed *sce1-4* with *tic40-4* and *hsp93-V-1*, two TIC-complex-associated mutants. These mutants are chlorotic, due to defects in protein import across the chloroplast inner membrane, and in this respect are highly similar to *ppi1* (Kovacheva et al., 2005). Significantly, the resulting double mutants, *tic40-4 sce1-4* and *hsp93-V-I sce1-4*, were indistinguishable from *tic40-4* and *hsp93-V-1*, their respective single mutant controls (Figures 2A and 2C). Moreover, the double mutants did not display changes in leaf chlorophyll accumulation relative to the single mutant controls (Figures 2B and 2D). We therefore concluded that the suppression effects observed in *ppi1 sce1* plants were background-specific and associated with the TOC complex.

**Figure 2.**
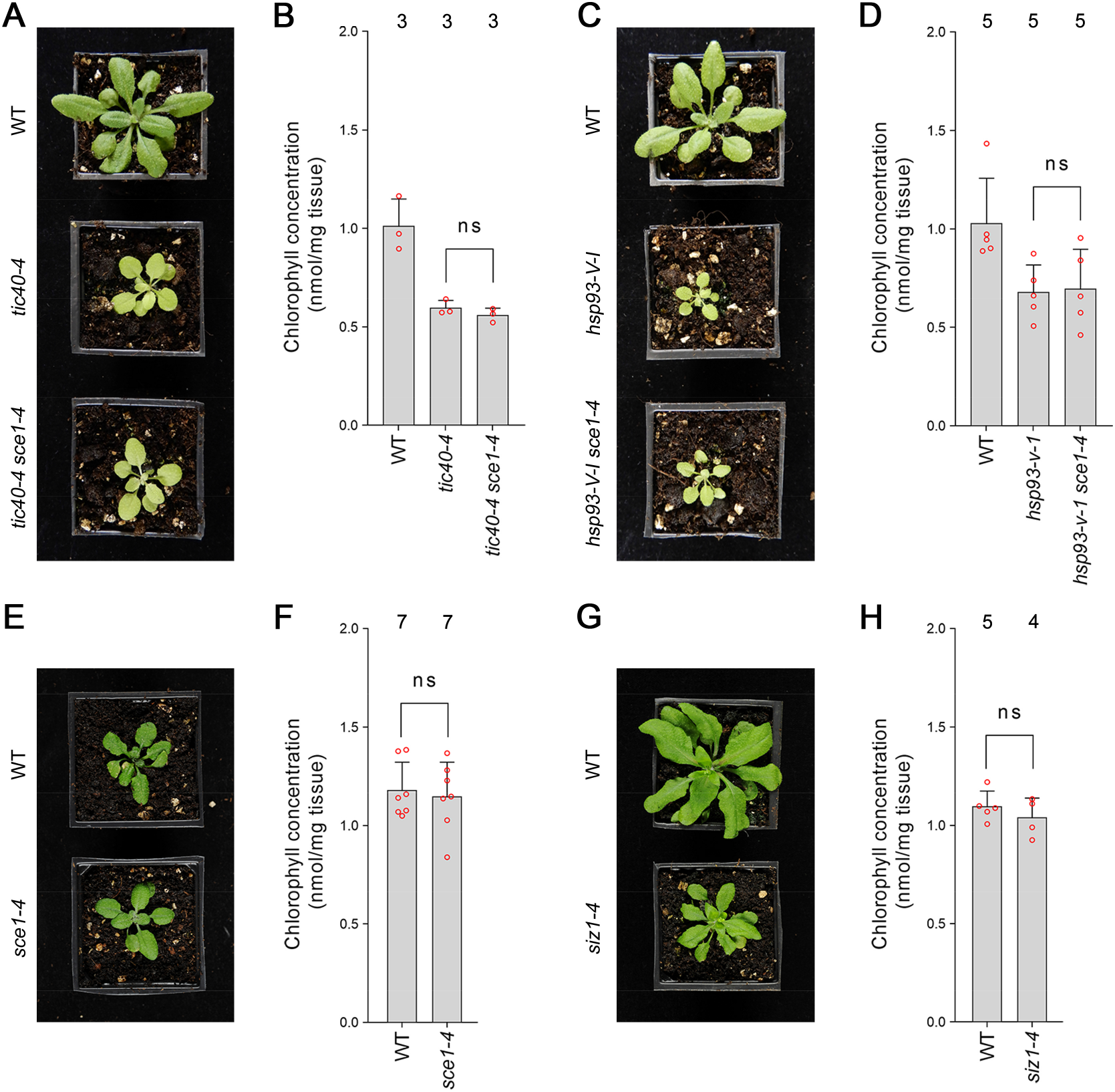
Genetic analysis reveals specificity of the suppression mediated by the *sce1-4* and *siz1-4* mutations. **(A)** The *tic40-4 sce1-4* double mutant did not appear greener than *tic40-4* after approximately four weeks of growth on soil. **(B)** The *tic40-4 sce1-4* double mutant did not show an enhanced accumulation of chlorophyll relative to *tic40-4* after approximately four weeks of growth on soil. Measurements were taken from the plants shown in (A) on the day of photography, as well as additional similar plants. There were no significant differences between the *tic40-4* and *tic40-4 sce1-4* plants (Two-tailed t-test, unpaired samples, T = 1.25, *p* = 0.280106). **(C)** The *hsp93-V-1 sce1-4* double mutant did not appear greener than *hsp93-V-1* after approximately four weeks of growth on soil. **(D)** The *hsp93-V-1 sce1-4* double mutant did not show an enhanced accumulation of chlorophyll relative to *hsp93-V-1* after approximately four weeks of growth on soil. Measurements were taken from the plants shown in (C) on the day of photography, as well as additional similar plants. There were no significant differences between the *hsp93-V-1* and *hsp93-V-1 sce1-4* plants (Two-tailed t-test, unpaired samples, T = 0.18, *p* = 0.860702). **(E)** The *sce1-4* single mutant did not appear greener than wild-type plants after approximately four weeks of growth on soil. **(F)** The *sce1-4* single mutant did not show an enhanced accumulation of chlorophyll relative to wild-type plants after approximately four weeks of growth on soil. Measurements were taken from the plants shown in (E) on the day of photography, as well as additional similar plants. There were no significant differences between the *sce1-4* and wild type plants (Two-tailed t-test, unpaired samples, T = 0.38, *p* = 0.708484). **(G)** The *siz1-4* single mutant did not appear greener than wild-type plants after approximately five weeks of growth on soil. **(H)** The *siz1-4* single mutant did not show an enhanced accumulation of chlorophyll relative to wild-type plants after approximately five weeks of growth on soil. Measurements were taken from the plants shown in (G) on the day of photography, as well as from additional similar plants. There were no significant differences between the *siz1-4* and wild type plants (T = 0.96, *p* = 0.370055). In all bar charts, error bars indicate standard deviation from the mean, and open red circles indicate individual data points. The numbers above the graphs indicate the number of biological replicates per sample. Statistical significance is indicated as follows: ns, not significant.

Next, we asked whether the *sce1-4* and *siz1-4* single mutants display an increase in chlorophyll concentration even in the wild-type background. However, neither mutant appeared greener than wild-type plants (Figures 2E and 2G; Figure 1 – Supplement 1A and 1C) or displayed an increase in leaf chlorophyll concentration (Figures 2F and 2H; Figure 1 – Supplement 1B and 1D). We therefore concluded that the suppression effects mediated by the SUMO mutants were synthetic phenotypes specific to the *ppi1* background.

### BiFC analysis reveals that SCE1 physically interacts with TOC proteins

Our reverse genetic experiments revealed a genetic interaction between the E2 SUMO conjugating enzyme, SCE1, and protein import across the chloroplast outer membrane. To determine whether SCE1 directly interacts with the TOC complex, we carried out bimolecular fluorescence complementation (BiFC) experiments in *Arabidopsis* protoplasts. The *SCE1* coding sequence was cloned into a vector that C-terminally appends the C-terminal half of YFP (cYFP) to its insert. This construct was co-expressed with various other constructs encoding TOC proteins bearing the complementary, N-terminal moiety of the YFP protein (nYFP), appended N-terminally; or with a negative control construct encoding ΔOEP7 bearing the nYFP moiety appended C-terminally. In this system, protein-protein interactions are inferred via the detection of a YFP signal, caused by the nYFP and cYFP fragments coming together to reconstitute a functional YFP protein.

Strikingly, SCE1-nYFP was found to physically associate with all tested TOC proteins – nYFP-Toc159, nYFP-Toc132, nYFP-Toc34 and nYFP-Toc33 (Figure 3). Moreover, these interactions were concentrated at the periphery of the chloroplasts, placing them in an appropriate subcellular context for the *in situ* regulation of the chloroplast protein import machinery. Conversely, SCE1-cYFP was not found to physically associate with the negative control protein ΔOEP7-nYFP. The ΔOEP7 protein comprises the transmembrane domain of plastid protein OEP7, which is sufficient to efficiently target the full-length YFP protein to the chloroplast outer membrane (Lee et al., 2001); thus cYFP-ΔOEP7 serves as a location-specific negative control.

**Figure 3.**
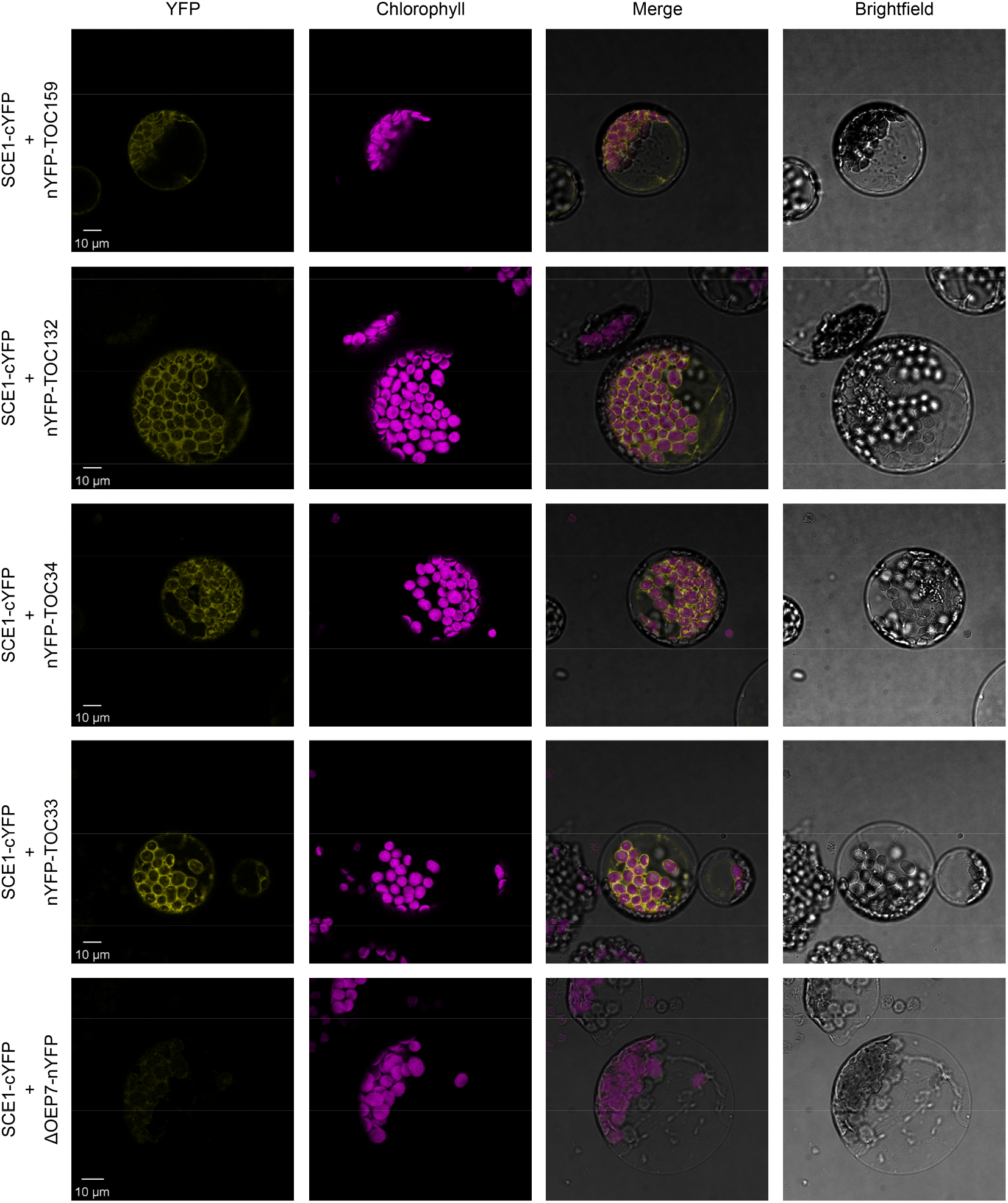
SCE1 physically interacts with TOC proteins *in vivo*. Bimolecular fluorescence complementation (BiFC) analysis of SCE1 protein-protein interactions was performed by imaging *Arabidopsis* protoplasts co-expressing proteins fused to complementary N-terminal (nYFP) and C-terminal (cYFP) fragments of the YFP protein, as indicated. Chlorophyll autofluorescence images were used to orientate the YFP signals in relation to the chloroplasts. SCE1 physically associated with all tested TOC proteins. The images shown are representative confocal micrographs indicating associations between SCE1 and Toc159, Toc132, Toc34 and Toc33. In contrast, SCE1 did not physically associate with ΔOEP7 (comprising the transmembrane domain of OEP7, which is sufficient to direct targeting to the chloroplast outer envelope), which served as a negative control. Scale bars = 10 μm.

### Manipulating the expression of three SUMO isoforms alters the phenotypic severity of *ppi1*

To further explore the genetic interaction between chloroplast protein import and the SUMO system, we crossed *ppi1* with several SUMO protein mutants. There are three major SUMO isoforms in *Arabidopsis* – SUMO1, SUMO2, and SUMO3. The *SUMO1* and *SUMO2* genes are expressed at a relatively high level throughout the plant and are largely functionally redundant (Saracco et al., 2007; van den Burg et al., 2010). In addition, they are highly similar to each other in terms of amino acid sequence (Saracco et al., 2007). In contrast, at steady state, *SUMO3* is expressed at a relatively low level throughout the plant, while the SUMO3 amino acid sequence is significantly divergent with respect to the other two SUMO isoforms (van den Burg et al., 2010).

First, we analysed *SUMO1* and *SUMO2*. We obtained *sum1-1* and *sum2-1*, two previously characterised null mutants (Saracco et al., 2007), and crossed them with *ppi1*. To account for the functional redundancy between these two genes, we also sought a *ppi1 sum1-1 sum2-1* triple mutant. However, as *SUMO1* and *SUMO2* are collectively essential, *ppi1 sum1-1 sum2-1* plants that were homozygous with respect to *ppi1* and *sum2-1*, but heterozygous with respect to the *sum1-1* mutation, were selected from a segregating population. The double and triple mutants were phenotypically characterised, and all three appeared larger and greener than the *ppi1* control plants (Figure 4A). Moreover, the double and triple mutants showed corresponding increases in leaf chlorophyll concentration, with the triple mutant showing a larger increase than the double mutants (Figure 4B). These were synthetic effects, as the *sum1-1*, *sum2-1*, and *sum1-1 sum2-1* single and double mutants did not appear greener than wild-type plants, or show increases in chlorophyll accumulation (Figure 4 – Supplement 1). We therefore concluded that the *sum1-1* and *sum2-1* mutants can additively suppress the phenotype of *ppi1*.

**Figure 4.**
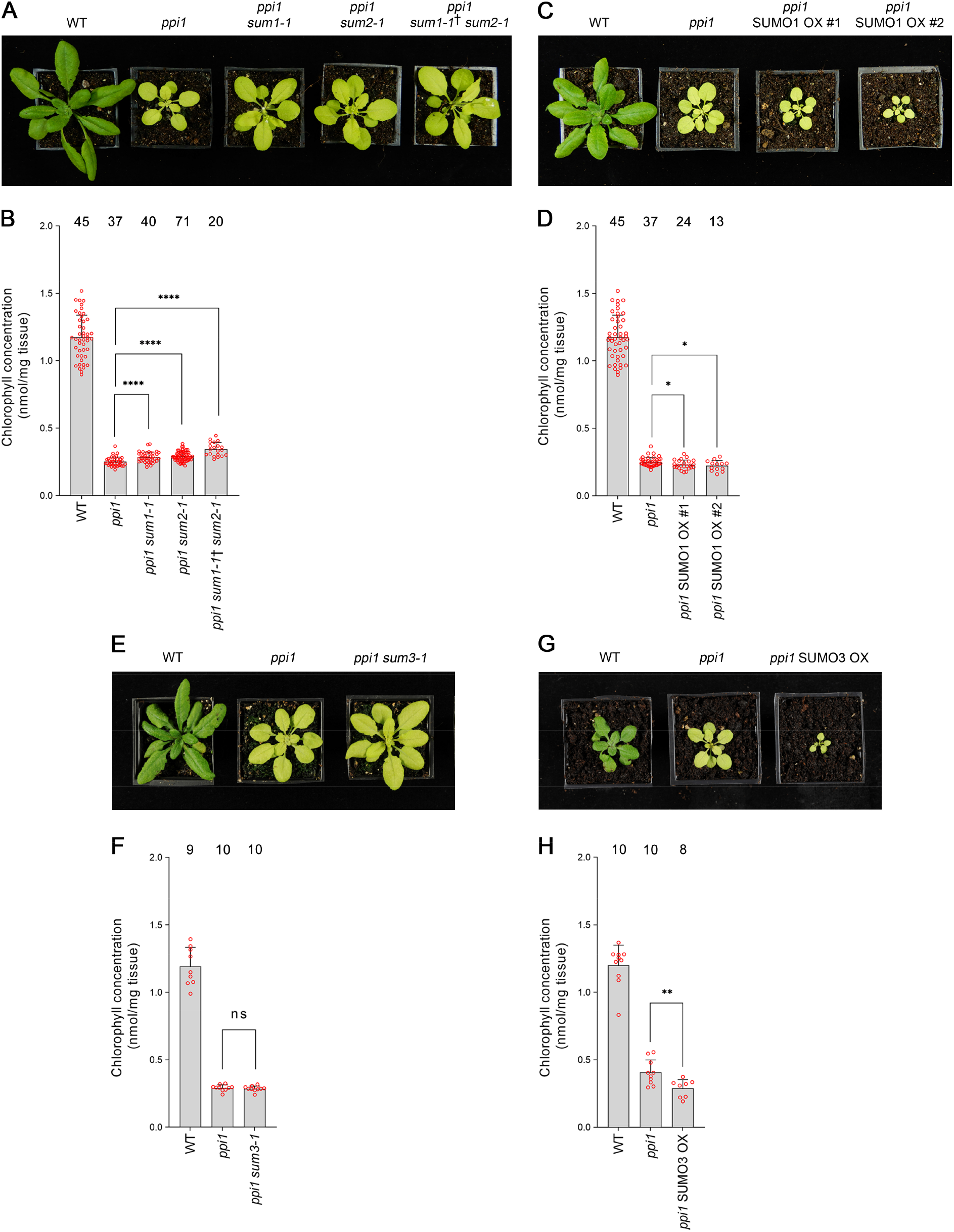
Genetic interactions between *ppi1* and the genes encoding three SUMO isoforms. **(A)** The *ppi1 sum1-1*, *ppi1 sum2-1*, and *ppi1 sum1-1*† *sum2-1* double and triple mutants appeared greener than *ppi1* after approximately four weeks of growth on soil. The dagger symbol indicates that the triple mutant was heterozygous with respect to the *sum1-1* mutation. **(B)** The *ppi1 sum1-1*, *ppi1 sum2-1*, and *ppi1 sum1-1*† *sum2-1* double and triple mutants showed enhanced accumulation of chlorophyll relative to *ppi1* after approximately four weeks of growth on soil. Measurements were taken from the plants shown in (A) on the day of photography, as well as additional similar plants. There were significant differences between the samples, as measured via a one-way ANOVA (F = 26.21, p = 6.65 × 10-14). A post-hoc Tukey HSD test indicated that there was a significant difference between the *ppi1* and *ppi1 sum1-1* samples (p < 0.00001). There were also significant differences between the *ppi1* and *ppi1 sum2-1* samples (p < 0.00001), and between the *ppi1* and *ppi1 sum1-1*† *sum2-1* samples (p < 0.00001). **(C)** The *ppi1* SUMO1 overexpression (OX) lines appeared smaller and paler than *ppi1* after approximately four weeks of growth on soil. **(D)** The *ppi1* SUMO1 OX lines showed reduced accumulation of chlorophyll relative to *ppi1* after approximately four weeks of growth on soil. Measurements were taken from the plants shown in (C) on the day of photography, as well as additional similar plants. There were significant differences between the *ppi1* and *ppi1* SUMO1 OX #1 plants (Two-tailed t-test, unpaired samples, T = 2.27, p = 0.026832), and between the *ppi1* and *ppi1* SUMO1 OX #2 plants (Two-tailed t-test, unpaired samples, T = 2.49, p = 0.01634). **(E)** The *ppi1 sum3-1* double mutant did not appear greener than *ppi1* after approximately four weeks of growth on soil. **(F)** The *ppi1 sum3-1* double mutant did not show an enhanced accumulation of chlorophyll relative to *ppi1* after approximately four weeks of growth on soil. Measurements were taken from the plants shown in (E) on the day of photography, as well as additional similar plants. There were no significant differences between the *ppi1* and *ppi1 sum3-1* plants (Two-tailed t-test, unpaired samples, T = 0.54, p = 0.59407). **(G)** The *ppi1* SUMO3 overexpression line appeared smaller and paler than *ppi1* after approximately three weeks of growth on soil. **(H)** The *ppi1* SUMO3 overexpression line showed reduced accumulation of chlorophyll relative to *ppi1* after approximately three weeks of growth on soil. Measurements were taken from the plants shown in (G) on the day of photography, as well as additional similar plants. There were significant differences between the *ppi1* SUMO3 OX plants and *ppi1* (Two-tailed t-test, unpaired samples, T = 2.99, p = 0.008688). In all bar charts, error bars indicate standard deviation from the mean, and open red circles indicate individual data points. The numbers above the graphs indicate the number of biological replicates per sample. Statistical significance is indicated as follows: ns, not significant; *, *p*<0.05; **, *p*<0.01; ****, *p*<0.0001.

To complement the preceding experiment, we generated transgenic plants overexpressing *SUMO1* in the *ppi1* background. The *SUMO1* coding sequence was cloned into a vector carrying a strong, constitutive promotor (cauliflower mosaic virus 35S) upstream of the cloning site. The resulting construct was stably introduced into the *ppi1* background via *Agrobacterium*-mediated transformation. Two lines carrying a single, homozygous transgene insert were identified and taken forward for analysis. The overexpression of *SUMO1* was confirmed in both lines by RT-PCR (Figure 4 – Supplement 2A). Significantly, both lines displayed an accentuation of the *ppi1* phenotype: the plants were significantly smaller and paler than the *ppi1* control plants (Figure 4C), and showed decreases in leaf chlorophyll concentration (Figure 4D).

Next, we turned our attention to *SUMO3*. We obtained *sum3-1*, a previously characterised null mutant (van den Burg et al., 2010), and crossed it with *ppi1*. The resulting double mutant was phenotypically characterised, but it did not appear obviously different from the *ppi1* control (Figure 4E). Correspondingly, it did not display any clear increase in leaf chlorophyll concentration relative to *ppi1* (Figure 4F). To complement this experiment, we generated transgenic plants overexpressing *SUMO3* in the *ppi1* background, using the approach described above, and a line carrying a single, homozygous insert was identified and taken forward for analysis. The overexpression of *SUMO3* was confirmed by RT-PCR (Figure 4 – Supplement 2B). Interestingly, the transgenic plants showed a striking increase in the severity of the *ppi1* phenotype: the plants were severely dwarfed and paler than the *ppi1* control (Figure 4G), and displayed a significant decrease in leaf chlorophyll accumulation (Figure 4H). These findings are particularly noteworthy when considered alongside a previous report which explored the consequences of overexpressing *SUMO3* in wild-type plants (van den Burg et al., 2010). In that study, *SUMO3* overexpression was not found to alter the appearance of the transgenic plants, which implies a degree of specificity in the phenotypic accentuation observed here.

### Biochemical analysis reveals SUMOylation of TOC proteins *in vivo*

The genetic and molecular experiments described thus far strongly suggested that TOC proteins are SUMOylated. However, to our knowledge, conclusive evidence that chloroplast-resident proteins are SUMOylated is currently lacking. To investigate whether chloroplast proteins may be SUMOylated, we isolated chloroplasts from seedlings by cell fractionation and analysed them by anti-SUMO immunoblotting. For this analysis, we employed a proven commercial antibody against SUMO1, which is one of the most abundant SUMO isoforms in *Arabidopsis* making it more tractable for analysis, and which furthermore is known to accumulate in response to heat and other stresses (Kurepa et al., 2003; van den Burg et al., 2010). To enhance detection of SUMOylated proteins in our samples, we subjected some of the seedlings to heat shock before chloroplast isolation and/or treatment with 10 mM N-ethylmaleimide (NEM) during chloroplast isolation. NEM is a potent inhibitor of SUMO-specific proteases (Hilgarth and Sarge, 2005). Importantly, we detected protein SUMOylation in the isolated chloroplast samples, and this SUMOylation was increased by NEM treatment (Figure 5 – Supplement 1).

Next, we carried out a series of biochemical experiments to determine whether TOC proteins are SUMOylated. In the first of these, the *SCE1* coding sequence was cloned into a vector that appends a C-terminal YFP tag (Karimi et al., 2002). We confirmed that the resulting SCE1-YFP construct delivers good expression and the expected nucleocytoplasmic fluorescence pattern when transiently expressed in protoplasts (Figure 5 – Supplement 2A). Then, we transfected a large volume of protoplasts with the SCE1-YFP construct (or with a YFP-HA negative control construct), and performed immunoprecipitation (IP) using YFP-Trap magnetic beads. The samples were analysed by immunoblotting. The YFP-HA and SCE1-YFP fusion proteins both showed robust expression and strong recovery in the IP elutions (Figure 5A). Remarkbly, the SCE1-YFP fusion protein was found to be associated with native Toc159 and Toc132, but not with the negative control proteins Tic110 or Tic40 (Figure 5A). Conversely, YFP-HA did not associate with any of the tested proteins. Given that SCE1 is a promiscuous enzyme that associates with thousands of proteins (Elrouby and Coupland, 2010), and that these interactions are likely to be transient, it is remarkable that TOC co-elution was detectable in this experiment.

**Figure 5.**
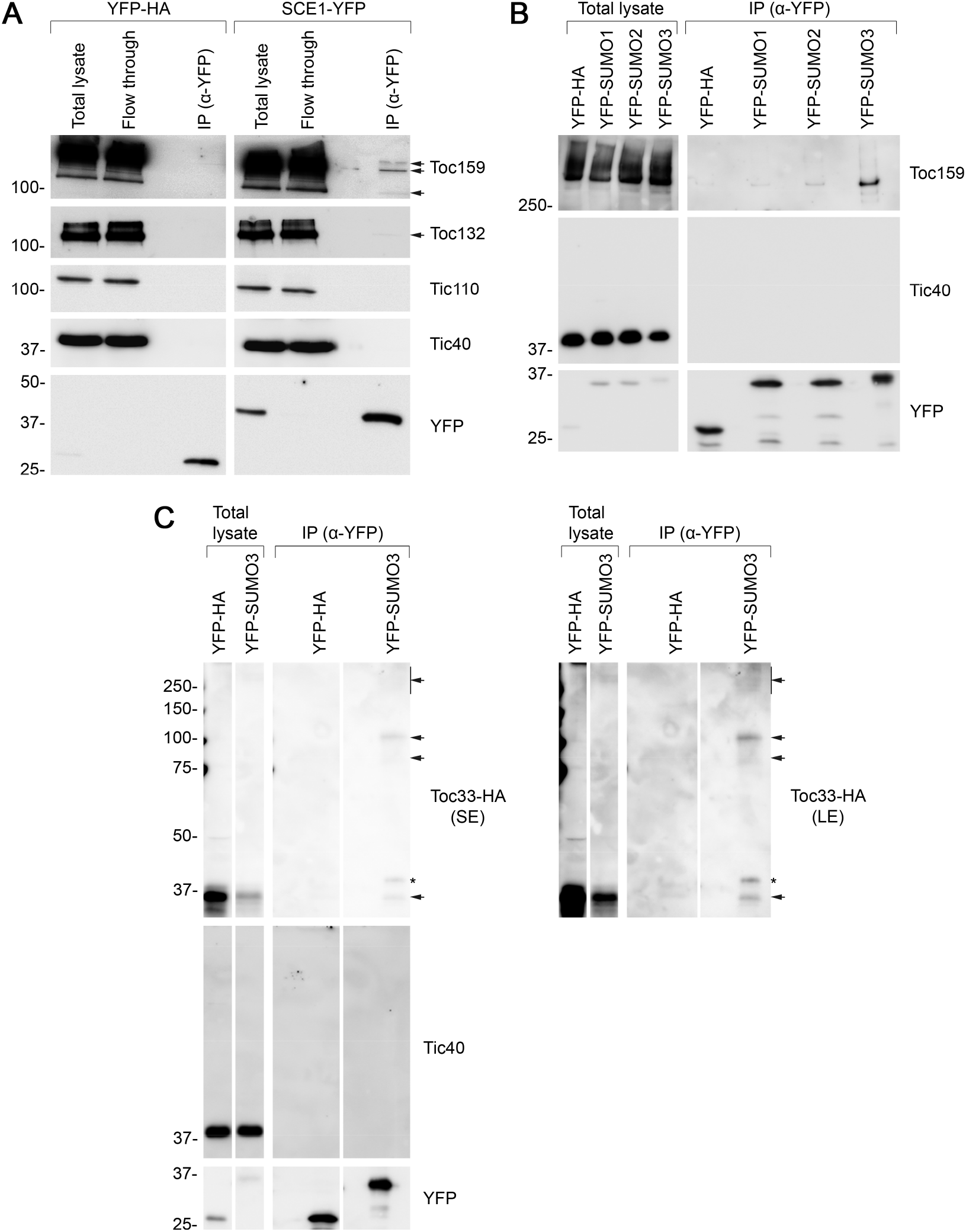
Immunoprecipitation analysis reveals that TOC proteins are SUMO targets. **(A)** SCE1 physically associated with native TOC proteins. *Arabidopsis* protoplasts expressing YFP-HA (left panel) or SCE1-YFP (right panel) were solubilised and subjected to immunoprecipitation (IP) analysis. The YFP-HA construct served as a negative control. In both cases, three samples were analysed: The ‘Total lysate’ sample (total protein extract from solubilised protoplasts); the ‘Flow through’ sample (the total protein sample after incubation with anti-YFP beads); and the ‘IP’ sample (the eluted fraction of the total protein sample that bound to the anti-YFP beads). The samples were analysed by immunoblotting, revealing that SCE1-YFP, but not the YFP-HA control, was associated with native Toc159 and Toc132 (indicated by the two arrows). Neither SCE1-YFP nor YFP-HA was associated with native Tic110 or Tic40, which were included as negative control proteins. **(B)** All three SUMO isoforms physically associated with native Toc159. Protoplasts expressing YFP-HA, YFP-SUMO1, YFP-SUMO2 or YFP-SUMO3 were solubilised and subjected to IP analysis as in (A). In all four cases, two samples (the ‘Total lysate’ and the ‘IP’ samples) were analysed by immunoblotting. Toc159 was resolved on an 8% acrylamide gel for four hours to maximise the resolution of high molecular weight bands. All three YFP-SUMO proteins were found to associate with native Toc159; however, YFP-SUMO3 immunoprecipitated Toc159 with the greatest efficiency. None of the four YFP fusion proteins associated with native Tic40, which served as a negative control protein. **(C)** YFP-SUMO3 physically associated with Toc33-HA and related high molecular weight species. Protoplasts co-expressing YFP-SUMO3 or YFP-HA together with Toc33-HA were solubilised and subjected to IP analysis as in (A). In both cases, two samples (the ‘Total lysate’ and ‘IP’ samples) were analysed by immunoblotting. The results showed that YFP-SUMO3, but not YFP-HA, was associated with Toc33-HA (indicated by the arrow). Bands corresponding to the molecular weight of Toc33-HA bearing one, two or several YFP-SUMO3 motifs were detected on the membrane (indicated by the upper arrows). The predicted molecular weight of YFP-SUMO3 is approximately 38.9 kDa. Neither YFP-SUMO3 nor YFP-HA was associated with Tic40, which served as a negative control protein. The asterisk indicates a nonspecific band. Short (SE) and long (LE) exposures are shown. Migration positions of standards are displayed to the left of the gel images, and sizes are indicated in kDa. The unprocessed membrane images are displayed in Source data 1.

In the second experiment, we cloned the *SUMO1*, *SUMO2* and *SUMO3* coding sequences into a vector that appends an N-terminal YFP tag to its insert (Karimi et al., 2002), a modification which previous studies have shown to be tolerated (Ayaydin and Dasso, 2004). All three constructs expressed well and showed the expected nucleocytoplasmic fluorescence pattern when transiently expressed in protoplasts (Figure 5 – Supplement 2B). The three constructs were expressed in parallel in protoplasts alongside the YFP-HA negative control construct. As in the previous experiment, the protoplasts were subjected to YFP-Trap immunoprecipitation, and the samples were subsequently analysed by immunoblotting. Remarkably, all three YFP-SUMO proteins were found to physically associate with Toc159, although YFP-SUMO3 clearly bound Toc159 with the greatest affinity (Figure 5B; Figure 5 – Supplement 3). Moreover, inspection of an extended exposure of the anti-YFP blot revealed a number of higher molecular weight bands that we interpret to be SUMO adducts and indicative of the functionality of the fusions (Figure 5 – Supplement 4). In contrast with the SUMO fusions, the YFP-HA negative control did not associate with Toc159, and none of the four YFP fusion proteins physically associated with Tic40, a negative control protein (Figure 5B).

The immunoprecipitation experiment described above identified SUMO3 as having the highest affinity for Toc159. To extend our analysis of SUMO3 to include another TOC proteins, and to provide more direct evidence for TOC protein SUMOylation, the experiment was repeated with modifications, as follows. Protoplasts were co-transfected with YFP-SUMO3 and Toc33-HA, or YFP-HA and Toc33-HA; in each case, Toc33 was transiently overexpressed to aid detection of this component and its adducts. Upon co-expression of these construct pairs, the protoplast samples were subjected to YFP-Trap immunoprecipitation analysis, as described earlier. In accordance with the Toc159 result (Figure 5B), YFP-SUMO3, but not YFP-HA, was found to physically associate with Toc33-HA (Figure 5C). Moreover, bands of the exact expected molecular weight for Toc33-HA bearing one or two YFP-SUMO3 moieties (75 and 114 kDa) were also detected. These bands were accompanied by a high molecular weight smear at the top of the immunoblot, which is indicative of complex, multisite or chain SUMOylation.

## Discussion

This work has revealed a genetic and molecular link between the SUMO system and chloroplast protein import. The genetic experiments demonstrated that SUMO system mutations can suppress the phenotype of the Toc33 mutant, *ppi1*, while the molecular and biochemical experiments indicated that TOC proteins associate with key SUMO system proteins and are SUMOylated. Visible suppression effects observed in the *ppi1* / SUMO system double mutants were linked to improvements in chloroplast development and enhanced accumulation of key TOC proteins. Thus, our results suggest that SUMOylation acts to destabilise the TOC complex, and that when such SUMOylation is perturbed the TOC proteins are stabilised. We interpret that TOC complexes containing Toc34, Toc75 and Toc159 accumulate at higher levels in *ppi1* / SUMO system double mutants, and that this synthetically improves the double mutant phenotypes relative to the *ppi1* control. Importantly, each core TOC protein, including all of those analysed in this study, was predicted with high probability to have one or more SUMOylation sites (Table 1) (Zhao et al., 2014; Beauclair et al., 2015).

**Table 1.** Bioinformatic analysis predicts that the core TOC proteins in *Arabidopsis* contain SUMOylation sites and SUMO interaction motifs. The GPS-SUMO algorithm was applied to the amino acid sequences of Toc159, Toc132, Toc120, Toc90, Toc75, Toc33 and Toc34 using the ‘high stringency’ setting, and the results generated are shown in columns 2, 3 and 4. ‘Consensus’ sites fall within canonical SUMO site motifs: Ψ-K-X-E (where Ψ indicates a hydrophobic amino acid, and X indicates any amino acid residue). ‘Non-consensus’ sites do not fall within canonical SUMO site motifs; analysis shows that ~40% of SUMOylation may occur at non-consensus sites (Zhao et al., 2014). ‘SUMO Interaction’ sites are predicted to mediate the non-covalent interaction between proteins and SUMO peptides. The JASSA algorithm was also applied to the amino acid sequences using the ‘high cut-off’ setting (see column 5). aa denotes amino acids.

| Protein name | Position (aa) | p-value | Type | Also predicted by JASSA (high stringency)? |
| --- | --- | --- | --- | --- |
| **atToc159**<br>(At4g02510)<br>1503 aa | 95 | 0.021 | Consensus | No |
|  | 106 | 0.034 | Consensus | Yes |
|  | 126 | 0.032 | Consensus | Yes |
|  | 144 | 0.02 | Consensus | Yes |
|  | 151 | 0.02 | Consensus | No |
|  | 246-250 | 0.005 | SUMO Interaction | Yes |
|  | 408-412 | 0.009 | SUMO Interaction | Yes |
|  | 486-490 | 0.009 | SUMO Interaction | Yes |
|  | 498 | 0.022 | Consensus | No |
|  | 502 | 0.049 | Non-consensus | No |
|  | 539 | 0.026 | Consensus | No |
|  | 1300 | 0.049 | Non-consensus | No |
|  | 1370 | 0.01 | Consensus | Yes |
| **atToc132**<br>(At2g16640)<br>1206 aa | 30 | 0.006 | Consensus | Yes |
|  | 66 | 0.036 | Consensus | Yes |
|  | 352 | 0.002 | Consensus | No |
|  | 895 | 0.005 | Consensus | No |
|  | 1077 | 0.014 | Consensus | No |
| **atToc120**<br>(At3g16620)<br>1084 aa | 52 | 0.008 | Consensus | No |
|  | 57 | 0.031 | Consensus | No |
|  | 209 | 0.05 | Non-consensus | No |
|  | 777 | 0.006 | Consensus | No |
|  | 959 | 0.013 | Consensus | No |
| **atToc90**<br>(At5g20300)<br>793 aa | 191 | 0.027 | Consensus | Yes |
|  | 481 | 0.017 | Consensus | No |
|  | 711 | 0.02 | Consensus | No |
|  | 786 | 0.042 | Non-consensus | Yes |
| **atToc75**<br>(At3g46740)<br>818 aa | 434 | 0.049 | Non-consensus | No |
|  | 513 | 0.029 | Consensus | No |
| **atToc33**<br>(At1g02280)<br>297 aa | 291 | 0.044 | Non-consensus | No |
| **atToc34**<br>(At5g05000)<br>313 aa | 290 | 0.026 | Consensus | No |
|  | 298 | 0.043 | Non-consensus | Yes |

The *ppi1* suppression effects described here are remarkably similar to those mediated by the *sp1* and *sp2* mutations (Ling et al., 2012; Ling et al., 2019). Like *sp1* and *sp2*, SUMO system mutations can partially suppress *ppi1* with respect to chlorophyll concentration, TOC protein accumulation, and chloroplast development. This similarity suggests that SUMOylation may regulate the activity of the CHLORAD pathway. This is an attractive hypothesis, as both SUMOylation and the CHLORAD pathway are activated by various forms of environmental stress (Kurepa et al., 2003; Ling and Jarvis, 2015; Ling et al., 2019). One possibility is that the SUMOylation of TOC proteins promotes their CHLORAD-mediated degradation. Indeed, as already noted, the ability to carry out SUMOylation is negatively correlated with the stability of TOC proteins in the context of the developed plants studied here. However, it should be kept in mind that SUMOylation can both promote and antagonise the effects of ubiquitination, in different situations (Desterro et al., 1998; Ahner et al., 2013; Liebelt and Vertegaal, 2016); and so our results do not preclude the possibility that SUMOylation may have different consequences for chloroplast biogenesis in other contexts. The Toc159 receptor is regulated by SP1 when integrated into the outer envelope membrane (Ling et al., 2012; Ling et al., 2019), but by a different E3 ligase when it exists as a cytosolic precursor during the earliest stages of development before germination (Shanmugabalaji et al., 2018). Thus, regulation by SUMOylation might be similarly different in these two distinct developmental contexts.

The precise mechanisms underpinning the observed negative regulation of the TOC apparatus by SUMOylation are currently unknown. One possibility is that the SUMOylation of TOC proteins promotes their association with SP1. SUMOylation can modify protein-protein interactions, and some RING-type E3 ubiquitin ligases specifically recognise SUMOylated substrates (Sriramachandran and Dohmen, 2014). However, these SUMO-targeted ubiquitin ligases (STUbLs) typically contain SUMO-interacting motifs (SIMs) which guide the ligases to SUMO proteins conjugated to their substrates, and these are not apparent in SP1 (data not shown) (Zhao et al., 2014). However, SP1 forms a complex with SP2 and very likely additional cofactors, and these could hypothetically provide a SUMO binding interface. Another possibility is that SUMOylation could be involved in the recruitment of Cdc48 from the cytosol. Two important Cdc48 cofactors are Ufd1 and Npl4, and the former contains a SUMO-interacting motif which can guide Cdc48 to SUMOylated proteins (Nie et al., 2012; Baek et al., 2013). Moreover, the SUMO-mediated recruitment of Cdc48 has important roles in the maintenance of genome stability in yeast (Bergink et al., 2013).

The biochemical experiments described in this manuscript indicate that, of the three SUMO isoforms tested, SUMO3 binds TOC proteins with the highest affinity. However, there is an apparent incongruence between the results of these experiments and the results of the genetic experiments. While the *sum1-1* and *sum2-1* mutants were found to additively suppress *ppi1*, the *sum3-1* mutant did not suppress *ppi1*. At face value, this seems puzzling; however, it can be explained by the relative abundance of the three SUMO proteins *in planta*. SUMO1 and SUMO2 are highly abundant relative to SUMO3, which is, at steady state, very weakly abundant (van den Burg et al., 2010). The immunoprecipitation data shown in Figure 5B indicated that SUMO1 and SUMO2 can weakly interact with Toc159, and so it is likely that these two isoforms can compensate for the loss of SUMO3 in the *sum3-1* mutant. Although SUMO3 associates with TOC proteins with the highest affinity, the higher abundance of the other two SUMO proteins may facilitate such compensation. It is also noteworthy that, when overexpressed, SUMO3 accentuates the *ppi1* phenotype to a far greater extent than does SUMO1.

It is now well established that the regulation of chloroplast protein import has critical roles in plant development and stress acclimation (Sowden et al., 2018; Watson et al., 2018). Here, we demonstrate regulatory crosstalk between the SUMO system and chloroplast protein import, and present results which are consistent with a model in which SUMOylation modulates the activity or effects of the CHLORAD pathway. The precise nature of the links between these two critically important control systems will be the subject of future investigation.

## Materials and methods

### Plant material and growth conditions

All *Arabidopsis thaliana* plants used in this work were of the Columbia-0 (Col-0) ecotype. The mutants used in most of the analyses (*ppi1*, *sce1-4*, *sum1-1*, *sum2-1*, *sum3-1*, *hsp93-V-I*, *tic40-4*) have been described previously (Jarvis et al., 1998; Kovacheva et al., 2005; Saracco et al., 2007; van den Burg et al., 2010). The *siz1-4* (SAIL_805_A10) and *siz1-5* (SALK_111280) mutants were obtained from the Salk Institute Genomic Analysis Laboratory (SIGnAL) (Alonso et al., 2003), via the Nottingham *Arabidopsis* Stock Centre (NASC). Each line was verified via PCR genotyping (see Table 2 for primer sequences) and phenotypic analysis (including the double and triple mutants). The positions of the T-DNA insertions were mapped via PCR. The following primer pairs were used to generate diagnostic amplicons from gDNA: LB1 and Siz1-Seq-1R (for mapping *siz1-4*), and LBb1 and Siz1-Seq-3R (for mapping *siz1-5*) (see Table 2D and 2E). The amplicons were sequenced and the positions of the T-DNA insertions inferred from the sequence data.

**Table 2.**
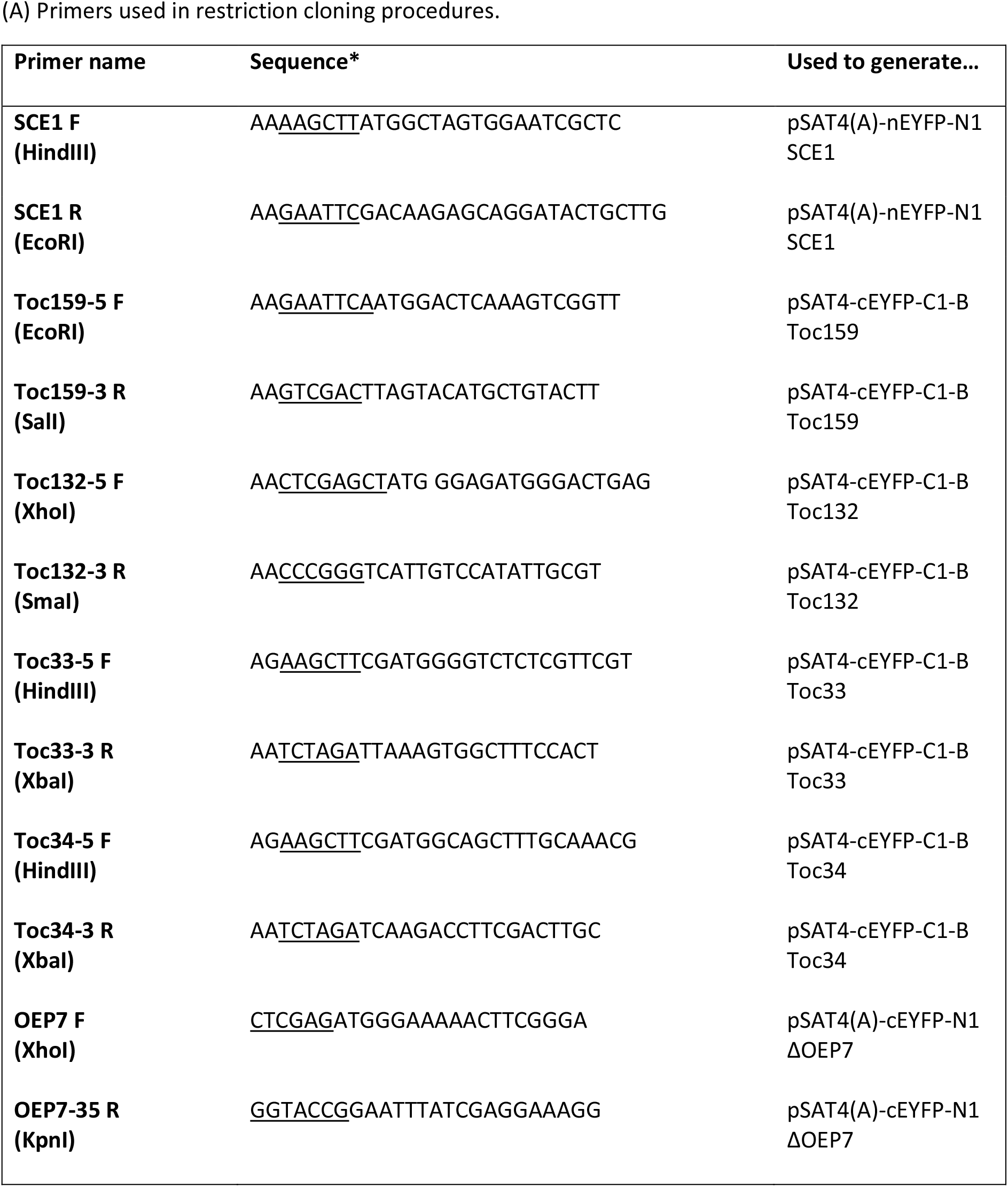
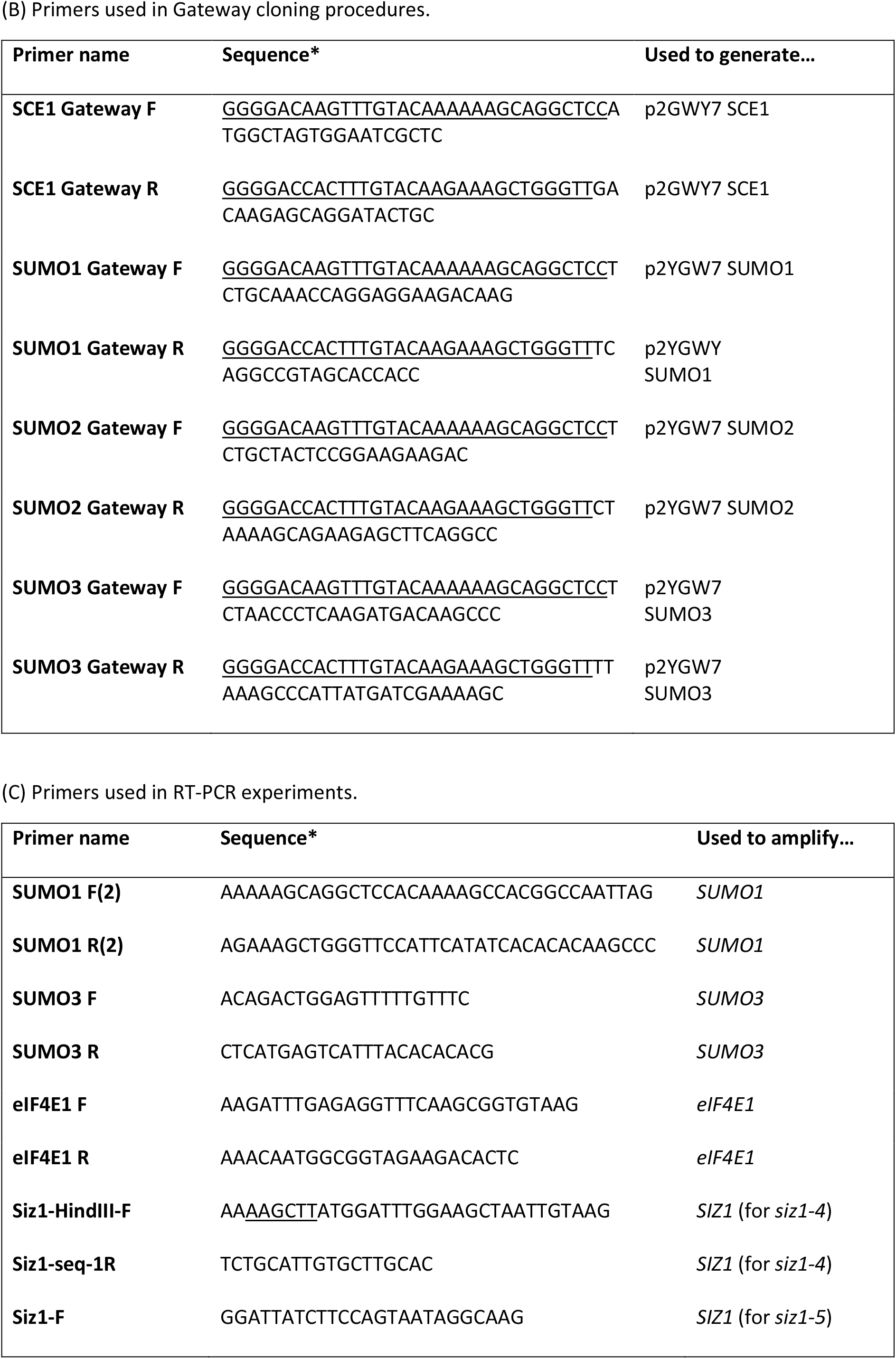
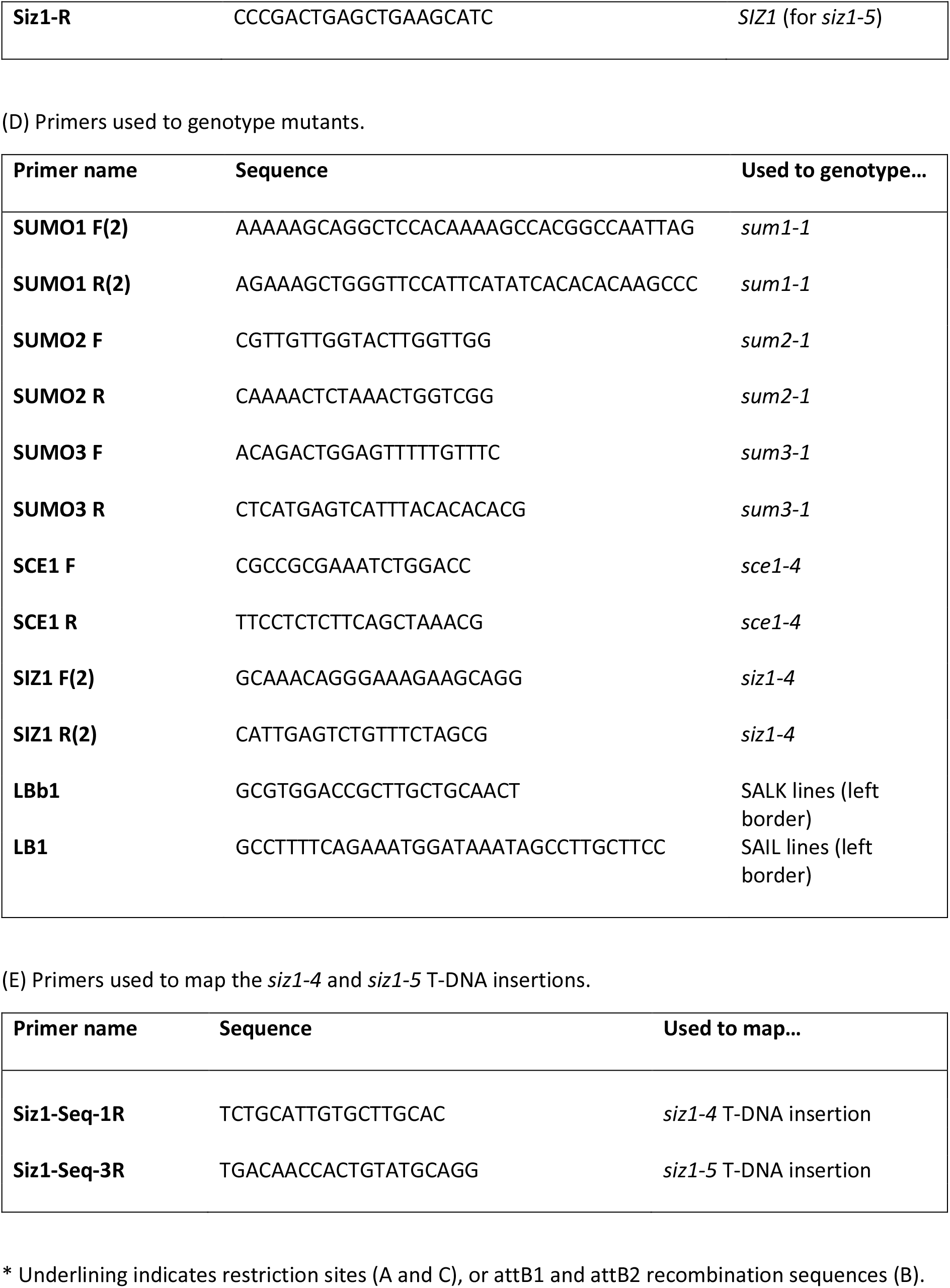
Primers used during the course of this study.

In most experiments, plants were grown on soil (80% (v/v) compost (Levington M2), 20% (v/v) vermiculite (Sinclair Pro, medium particle size)). However, where plants were grown for selection of transformants or for chloroplast isolation, seeds were surface sterilised and sown on petri plates containing Murashige-Skoog (MS) agar medium. The plates were stored at 4°C for 48 hours before being transferred to a growth chamber. Both soil-grown and plate-grown plants were kept in a growth chamber (Percival Scientific) under long-day conditions (16 hours light, 8 hours dark). The light intensity was approximately 120 μE m^−2^ s^−1^, the temperature was held constant at 20°C, and the humidity was held constant at approximately 70% (relative humidity).

### Chlorophyll measurements

Chlorophyll measurements were taken from mature rosette leaves in each instance. A handheld Konica-Minolta SPAD-502 meter was used to take each measurement, and the raw values were converted into chlorophyll concentration values (nmol/mg tissue) via published calibration equations (Ling et al., 2011).

### Chloroplast isolation and protein extraction

Chloroplasts were isolated from 14-day-old, plate-grown seedlings as described previously (Flores-Pérez and Jarvis, 2017). Some of the seedlings were heat-shocked immediately prior to chloroplast isolation. To do this, the plates containing the seedlings were wrapped in clingfilm and placed into a water bath (42°C for one hour, followed by a one hour recovery period at 22°C). Protein samples were prepared from the isolated chloroplasts by extraction using SDS-PAGE sample buffer, as well as from whole 14-day-old seedlings as previously described (Kovacheva et al., 2005). In some cases, the samples were treated with 10 mM N-ethylmaleimide (Hilgarth and Sarge, 2005); this was added directly to the protein extraction buffer (whole seedling samples), or to the chloroplast isolation buffer following polytron homogenization and all subsequent buffers (chloroplast samples).

### Plasmid constructs

The constructs used in the BiFC experiments were generated as follows. The coding sequences of *SCE1*, *SIZ1*, *TOC159*, *TOC132*, *TOC34* and *TOC33* were PCR amplified from wild-type cDNA (see Table 2 for primer sequences). In the case of *ΔOEP7*, the first 105 base pairs of the *OEP7* coding sequence were amplified; this encodes a truncated sequence which is sufficient to efficiently target the full-length YFP protein to the chloroplast outer envelope membrane (Lee et al., 2001). The inserts were cloned into one of the following complementary vectors: pSAT4(A)-cEYFP-N1 (*SCE1*), pSAT4-nEYFP-C1 (*TOC159*, *TOC132*, *TOC34*, *TOC33*), or pSAT4(A)-nEYFP-N1 (*ΔOEP7*), which were described previously (Tzfira et al., 2005; Citovsky et al., 2006).

The constructs used in the immunoprecipitation experiments were generated as follows. The coding sequences of *SCE1*, *SUMO1*, *SUMO2* and *SUMO3* were PCR amplified from wild-type cDNA using primers bearing 5’ attB1 and attB2 adaptor sequences (see Table 2 for primer sequences). The amplicons were then cloned into pDONR221 (Invitrogen), a Gateway entry vector. The inserts from the resulting entry clones were then transferred to one of two destination vectors: p2GWY7 (*SCE1*) or p2YGW7 (*SUMO1*, *SUMO2*, *SUMO3*); the former appends a C-terminal YFP tag to its insert, and the latter appends an N-terminal YFP tag to its insert (Karimi et al., 2002; Karimi et al., 2005). The Toc33-HA and YFP-HA constructs have been described previously (Ling et al., 2019).

The constructs used to generate transgenic plants were generated as follows. The coding sequences of *SUMO1* and *SUMO3* were PCR amplified from wild-type cDNA using primers bearing 5’ attB1 and attB2 adaptor sequences (see Table 2 for primer sequences). The inserts were then cloned into pDONR201 (Invitrogen), a Gateway entry vector. The inserts from the resulting entry clones were then transferred to the pH2GW7 binary destination vector (Karimi et al., 2002; Karimi et al., 2005).

### Transient expression assays

Protoplasts were isolated from mature rosette leaves of wild-type *Arabidopsis* plants and transfected in accordance with an established method (Wu et al., 2009; Ling et al., 2012). In the BiFC experiments, 100 μL protoplast suspension (containing approximately 10^5^ protoplasts) was transfected with 5 μg plasmid DNA; in the immunoprecipitation experiments, 600 μL protoplast suspension (containing approximately 6 × 10^5^ protoplasts) was transfected with 30 μg plasmid DNA. In both cases, the samples were analysed after 15-18 hours.

### Stable plant transformation

Transgenic lines carrying the *SUMO1-OX* or *SUMO3-OX* constructs were generated via *Agrobacterium*-mediated floral dip transformation (Clough and Bent, 1998). Transformed plants (T_1_ generation) were selected on MS medium containing phosphinothricin. Multiple T_2_ families were analysed in each case, and lines bearing a single T-DNA insertion were taken forward for further analysis. Transgene expression was analysed by semi-quantitative RT-PCR as described previously (Kasmati et al., 2011) (see Table 2 for primer sequences).

### Transmission electron microscopy

Transmission electron micrographs were recorded using mature rosette leaves as previously described (Huang et al., 2011). Images were taken from three biological replicates (different leaves from different individual plants), and at least 10 images were taken per replicate. The images were analysed using ImageJ (Schneider et al., 2012). The freehand tool was used to measure the plan area of the chloroplasts. For this, between 9 and 28 chloroplasts were analysed for each biological replicate (i.e., for each plant), and then an average value for each replicate was calculated and used for statistical comparisons. The analysis of chloroplast ultrastructure was performed as in previous work (Huang et al., 2011). For this, between 3 and 8 chloroplasts were analysed per biological replicate, and the data were processed as above.

### BiFC experiments

The BiFC experiments were carried out as described previously (Ling et al., 2019). Protoplasts were co-transfected with two constructs encoding fusion proteins bearing complementary fragments of the YFP protein (nYFP and cYFP) (Citovsky et al., 2006). After transfection and overnight incubation, the protoplasts were imaged using a Leica TCS SP5 laser scanning confocal microscope equipped with a Leica HC Plan Apochromat CS2 63.0x UV water immersion lens with a numerical aperture (N.A.) of 1.2. YFP was excited with an Argon-ion laser at 514 nm, selected using an acousto-optic tuneable filter (AOTF), and was detected using a 525-600 nm bandpass filter and a photomultiplier. Chlorophyll fluorescence was simultaneously excited with 514 nm excitation and detected with a 680-700 nm bandpass filter using a photomultiplier. Images were collected in 8-bit resolution with the pinhole set at 111.5 μm (1 Airy Unit), using 16-line averaging and a scan speed of 400 Hz. The image size was 512 × 512 pixels, with an (x,y) pixel size of 0.239 μm. Images were processed in the Leica Application Suite (LAS) software.

### Immunoblotting and immunoprecipitation

Protein extraction and immunoblotting were performed as described previously (Kovacheva et al., 2005). Total protein samples were extracted from 50 mg of intact, pooled seedlings after two weeks of growth. To detect proteins, we used an anti-SUMO1 antibody (Ab5316, Abcam), an anti-Toc75-III antibody (Kasmati et al., 2011), an anti-Toc159 antibody (Bauer et al., 2000), an anti-Toc132 antibody (Ling et al., 2012), an anti-Toc33 antibody (Kasmati et al., 2011), an anti-Tic110 antibody (Aronsson et al., 2010), an anti-Tic40 antibody (Kasmati et al., 2011), and an anti-green fluorescent protein antibody (Sigma, SAB4301138). In most cases, the secondary antibody used was goat anti-rabbit immunoglobulin G (IgG) conjugated with horseradish peroxidase (Sigma, 12-348); and protein bands were visualised via chemiluminescence using an ECL Plus Western blotting detection kit (GE Healthcare) and an LAS-4000 imager (Fujifilm). However, in the case of Figure 5 – Supplement 1, the secondary antibody was goat anti-rabbit IgG conjugated with alkaline phosphatase (Sigma, A3687), and the membrane was incubated with BCIP/NBT chromogenic substrate (Sigma, B3679).

The immunoprecipitation (IP) experiments were carried out as described previously (Ling et al., 2019). Constructs encoding YFP-conjugated fusion proteins (YFP-HA, SCE1-YFP, YFP-SUMO1, YFP-SUMO2, YFP-SUMO3) were transiently expressed in protoplasts. In some cases, the constructs were co-expressed with a construct encoding Toc33-HA. The protoplasts were solubilised using IP buffer containing 1% Triton X-100, and the resulting lysates were incubated with GFP-Trap beads (Chromotek). After four washes in IP buffer, the protein samples were eluted by boiling in SDS-PAGE loading buffer, and then analysed by immunoblotting.

### Statistical analysis

The data from each experiment were analysed in R. In most cases, two-tailed T-tests were performed. However, in one case, a one-way ANOVA was performed in conjunction with a Tukey HSD test (as indicated in the figure legend). The figures are annotated to indicate the level of significance, as follows: ns, not significant; *, *p*<0.05; **, *p*<0.01; ***, *p*<0.001; ****, *p*<0.0001; *****, *p*<0.00001.

### SUMO site prediction

The amino acid sequences of Toc159, Toc132, Toc120, Toc90, Toc75, Toc33 and Toc34 were retrieved from The *Arabidopsis* Information Resource (TAIR) website (Berardini et al., 2015). The GPS-SUMO algorithm was applied to all seven sequences (http://sumosp.biocuckoo.org/online.php) (Zhao et al., 2014). The ‘high stringency’ setting was applied. The *p*-values were generated by the GPS-SUMO algorithm, and hits that were accompanied by *p*-values exceeding *p>*0.05 were manually removed. The JASSA algorithm was also applied to all seven amino acid sequences (http://www.jassa.fr/index.php) (Beauclair et al., 2015). In this case, the ‘high cut-off’ setting was applied. The GPS-SUMO and JASSA algorithms use fundamentally different methodologies (Chang et al., 2018).

## Acknowledgements

We are very grateful to Alistair Haslam for initiating the genetic experiments. We also thank Dr Errin Johnson and Raman Dhaliwal (Dunn School of Pathology, University of Oxford) for assistance with the electron microscopy, and Professor Jane Langdale for advice during the course of the work. We are grateful to Pedro Bota and Rita Ross for technical support. This work was supported by the Oxford Interdisciplinary Bioscience Doctoral Training Partnership (DTP), and by grants from the Biotechnology and Biological Sciences Research Council (BBSRC) (grant numbers BB/K018442/1, BB/N006372/1, BB/R016984/1 and BB/R009333/1) to RPJ.

---

**Figure 1 – Supplement 1.**
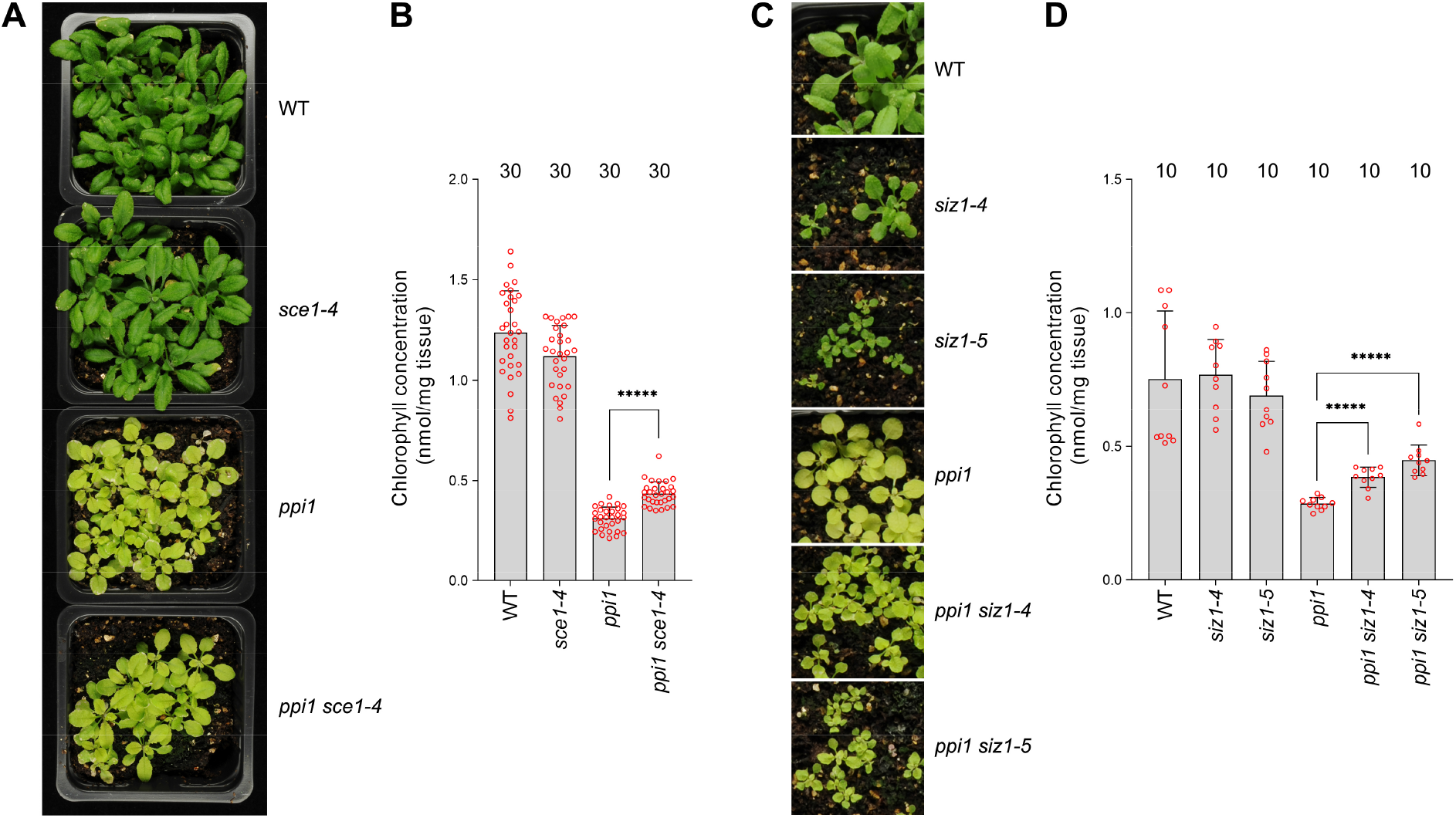
The E2 SUMO conjugating enzyme mutation, *sce1-4*, and the E3 SUMO ligase mutations, *siz1-4* and *siz1-5*, suppress the phenotype of plastid protein import mutant, *ppi1*. **(A)** The *ppi1 sce1-4* double mutant appeared greener than *ppi1* after approximately three weeks of growth on soil, while the single *sce1-4* mutant appeared no greener than the wild-type control. **(B)** The *ppi1 sce1-4* double mutant showed enhanced accumulation of chlorophyll relative to *ppi1* after approximately three weeks of growth on soil (T = 8.19, *p* < 0.0001), while the single *sce1-4* mutant displayed no enhanced accumulation of chlorophyll relative to the wild-type control. Measurements were taken from the plants shown in (A) on the day of photography, as well as additional similar plants. **(C)** The *ppi1 siz1-4* and *ppi1 siz1-5* double mutants appeared greener than *ppi1* after approximately two weeks of growth on soil, while the single *siz1-4* and *siz1-5* mutants appeared no greener than the wild-type control. **(D)** The *ppi1 siz1-4* and *ppi1 siz1-5* double mutants showed enhanced accumulation of chlorophyll relative to *ppi1* after approximately two weeks of growth on soil (T = 7.16, *p* < 0.0001 and T = 8.26, *p* < 0.0001 respectively), while the single *siz1-4* and *siz1-5* mutants displayed no enhanced accumulation of chlorophyll relative to the wild-type control. Measurements were taken from the plants shown in (C) on the day of photography, as well as additional similar plants. In all bar charts, error bars indicate standard deviation from the mean, and open red circles indicate individual data points. The numbers above the graphs indicate the number of biological replicates per sample. Statistical significance is indicated as follows: *****, *p*<0.00001.

**Figure 1 – Supplement 2.**
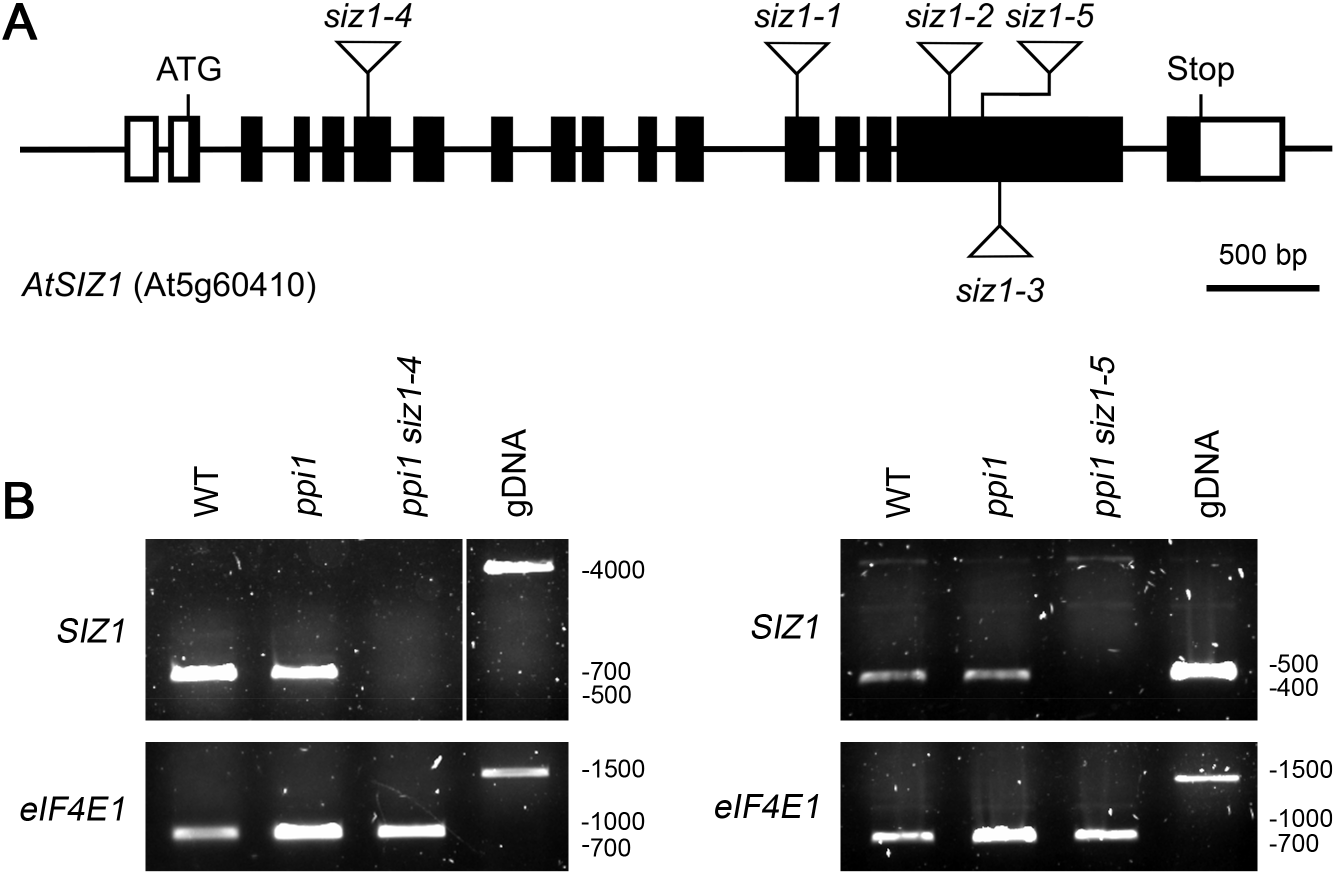
Molecular analysis of the *siz1-4* and *siz1-5* mutants. **(A)** Schematic representation of the *SIZ1* (At5g60410) gene, annotated with the positions of the *siz1* T-DNA insertions. The *siz1-4* and *siz1-5* insertion sites were verified in this study by DNA sequencing. The black boxes indicate exons, while the interconnecting black lines represent introns. The white boxes indicate untranslated regions. **(B)** Analysis of *SIZ1* mRNA expression in *siz1-4* and *siz1-5*. RNA samples were taken from two-week-old seedlings and used as a template for cDNA synthesis. Primers specific to *SIZ1* and the reference gene *eIF4E1* were used in two parallel RT-PCR experiments. gDNA, genomic DNA. Migration positions of standards are displayed to the right of the gel images, and sizes are indicated in base pairs.

**Figure 1 – Supplement 3.**
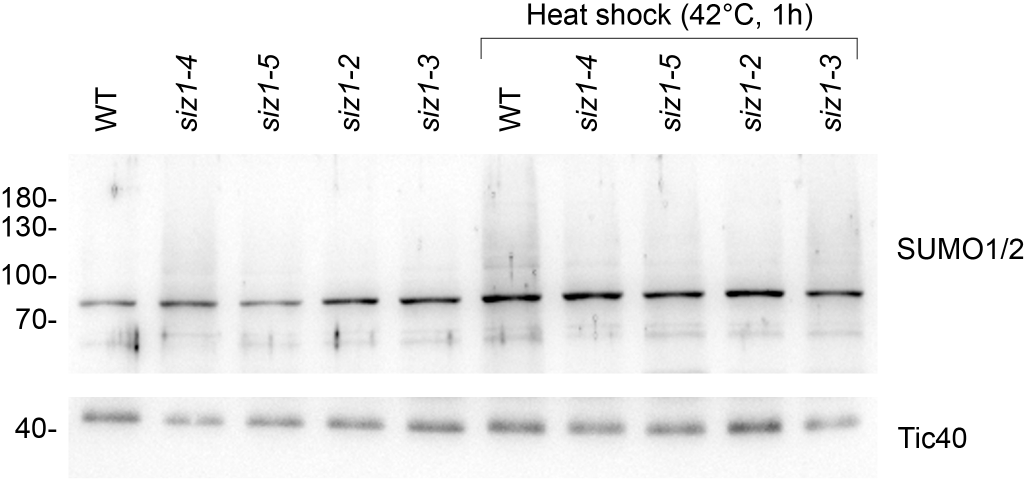
The *siz1-4* and *siz1-5* mutants display reduced global SUMOylation in response to heat shock. 14-day-old seedlings of the indicated genotypes were subjected to heat shock (42°C for 1 hour, followed by a 1-hour recovery period at 22°C). Protein samples were taken from whole seedlings and analysed by immunoblotting. The *siz1-4* and *siz1-5* mutants displayed a moderate reduction of global SUMOylation, and this reduction was similar in magnitude to the reduction observed in the previously characterised *siz1-2* and *siz1-3* mutants. The unprocessed membrane images are displayed in Source data 1.

**Figure 4 – Supplement 1.**
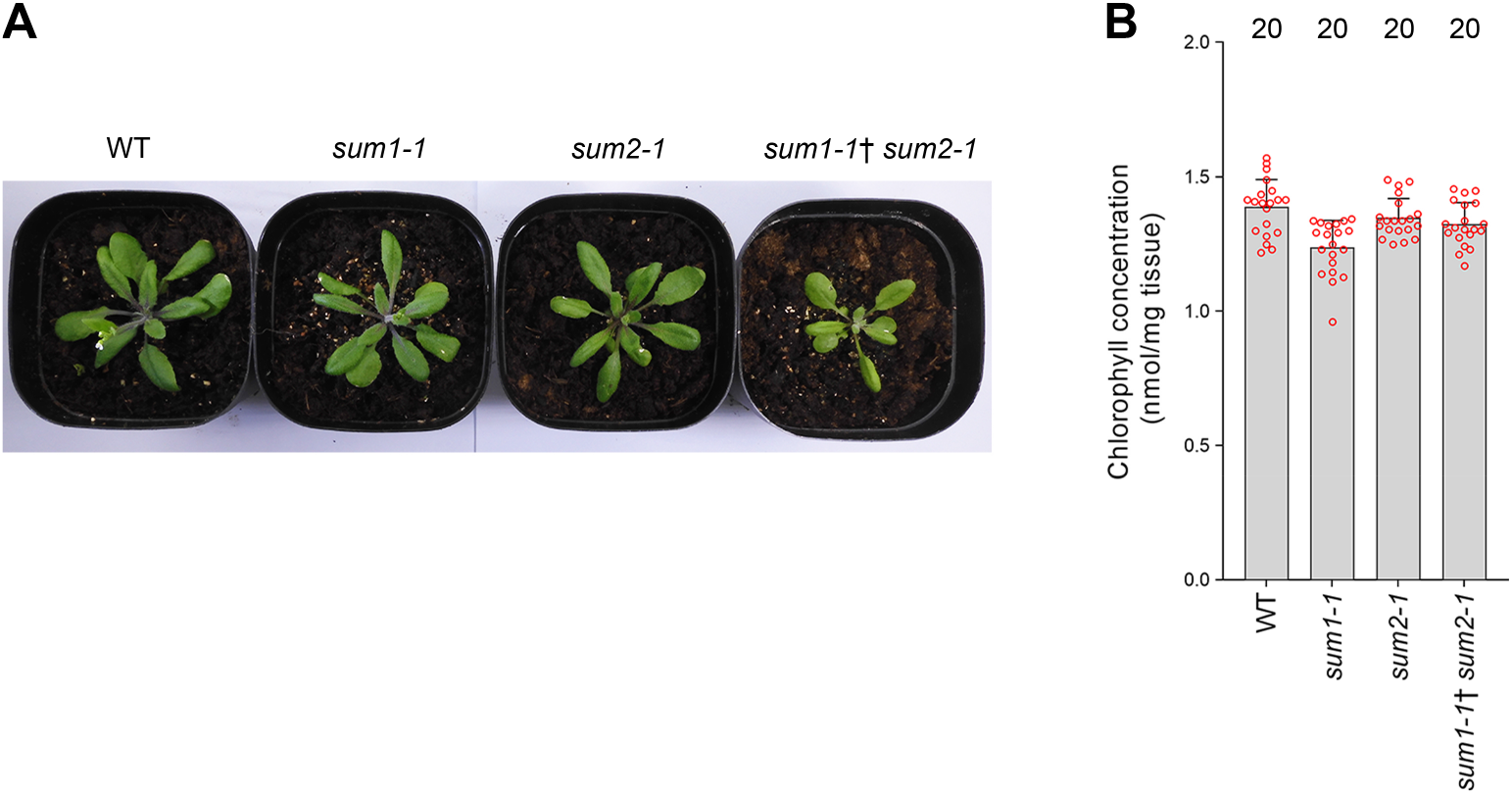
The *sum1-1*, *sum2-1*, and *sum1-1*^†^ *sum2-1* single and double mutants do not show enhanced chlorophyll accumulation relative to wild-type plants. **(A)** The *sum1-1*, *sum2-1*, and *sum1-1*^†^ *sum2-1* mutants did not appear greener than wild-type plants after approximately four weeks of growth on soil. **(B)** The *sum1-1*, *sum2-1*, and *sum1-1*^†^ *sum2-1* mutants did not show enhanced chlorophyll accumulation relative to wild-type plants after approximately four weeks of growth on soil. The dagger symbol indicates that the double mutant was heterozygous with respect to the *sum1-1* mutation. Error bars indicate standard deviation from the mean, and open red circles indicate individual data points. The numbers above the graph indicate the number of biological replicates per sample.

**Figure 4 – Supplement 2.**
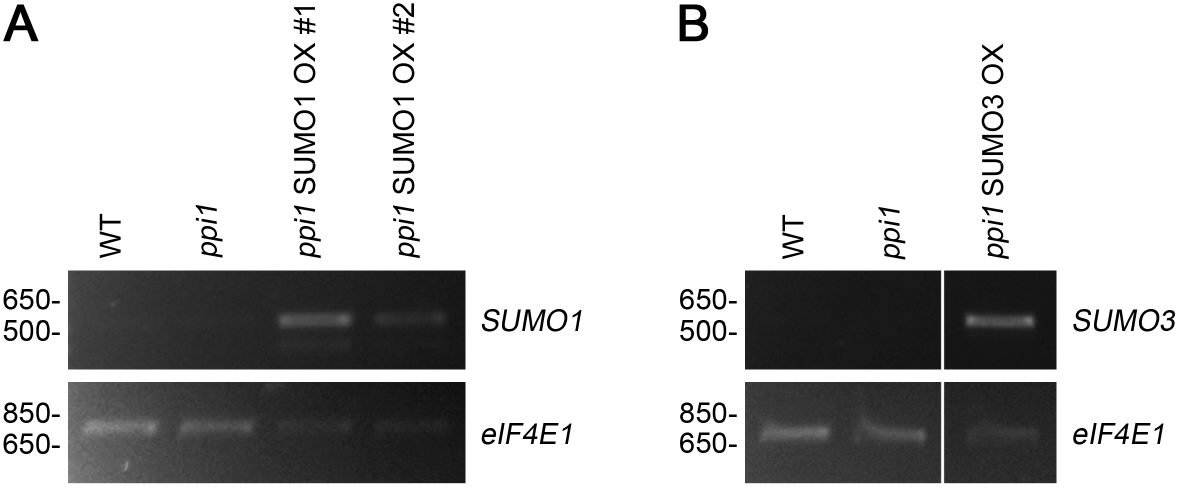
Analysis of the expression of the 35S:*SUMO1* and 35S:*SUMO3* transgenes in the selected transformants by RT-PCR. Total RNA was extracted from the rosette tissue of plants that had been grown on soil for approximately four weeks. The expression of *SUMO1* **(A)** or *SUMO3* **(B)**, and of the control gene, *eIF4E1*, was analysed by semi-quantitative RT-PCR. A limited number of amplification cycles (*n* = 26) were employed to prevent saturation. Migration positions of standards are displayed to the left of the gel images, and sizes are indicated in base pairs.

**Figure 5 – Supplement 1.**
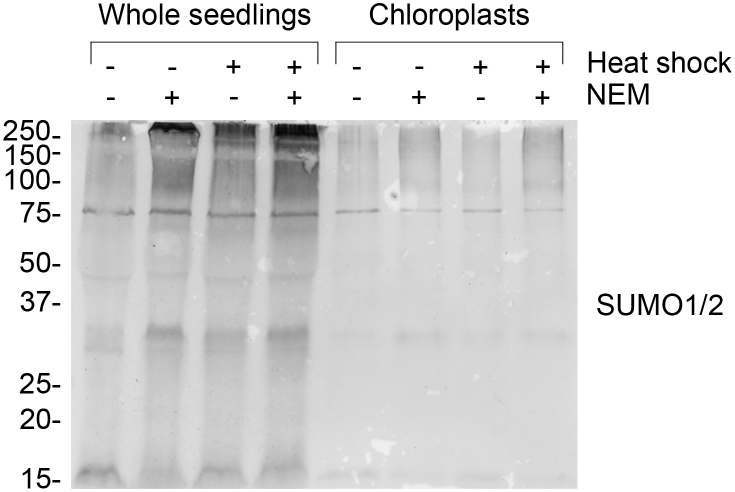
Chloroplast resident proteins are SUMOylated. Anti-SUMO1 immunoblot of protein samples taken from whole seedlings (left hand side) and isolated chloroplasts (right hand side). Where indicated, seedlings were exposed to heat shock (42°C for one hour) and/or incubated with 10 mM N-ethylmaleimide (NEM) to aid the detection of SUMOylated proteins. The anti-SUMO1 antibody shows significant cross-reactivity with SUMO2. Migration positions of standards are displayed to the left of the gel images, and sizes are indicated in kDa.

**Figure 5 – Supplement 2.**
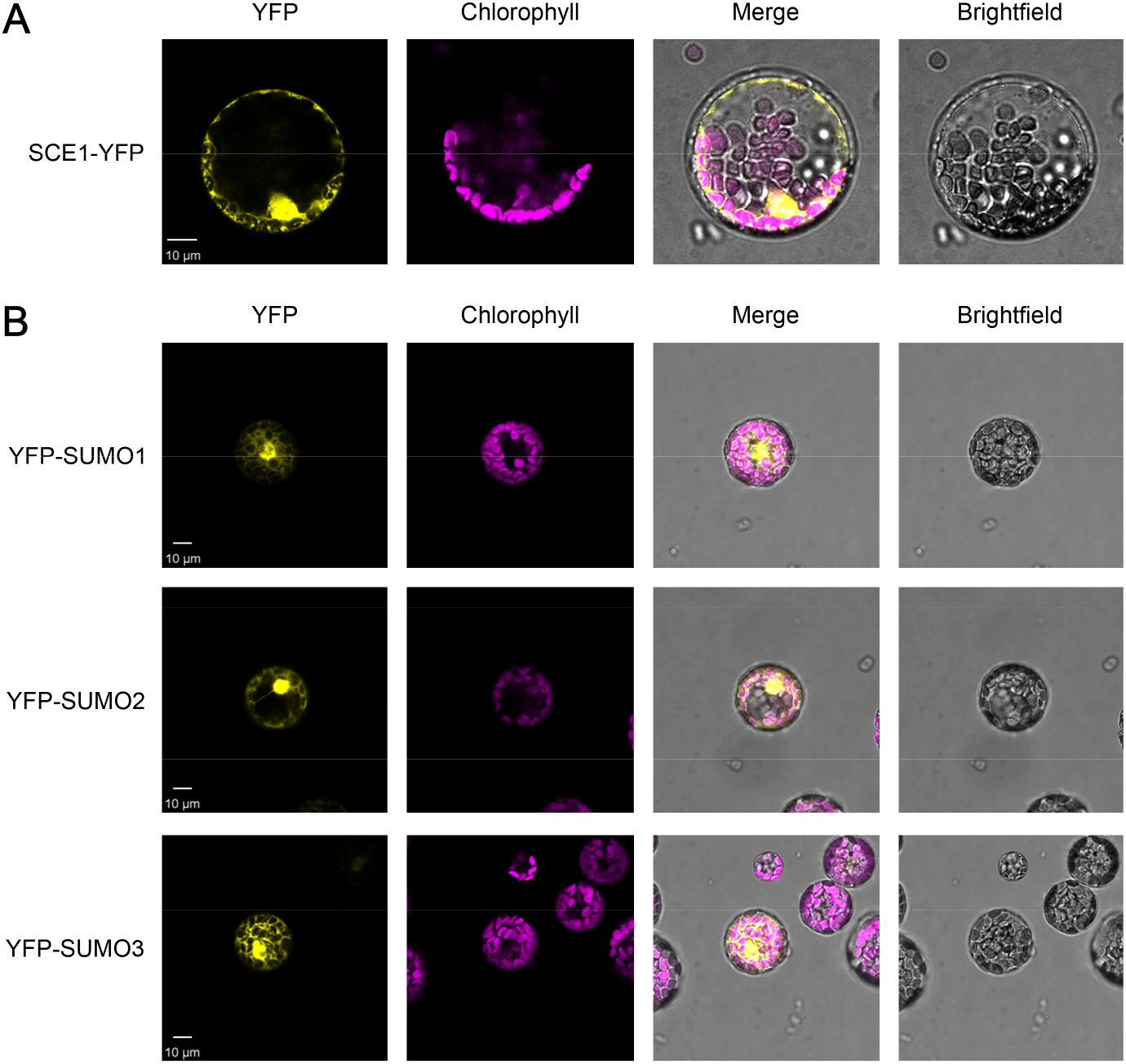
Analysis of the expression of the YFP-tagged constructs used in the immunoprecipitation experiments by confocal microscopy. The expression of the SCE1-YFP construct **(A)**, and of the three YFP-SUMO constructs **(B)**, was analysed and confirmed by imaging transfected *Arabidopsis* protoplasts. Chlorophyll autofluorescence images were employed to orientate the YFP signals in relation to the chloroplasts. Representative confocal micrographs show a typical protoplast in each case. Scale bars = 10 μm.

**Figure 5 – Supplement 3.**
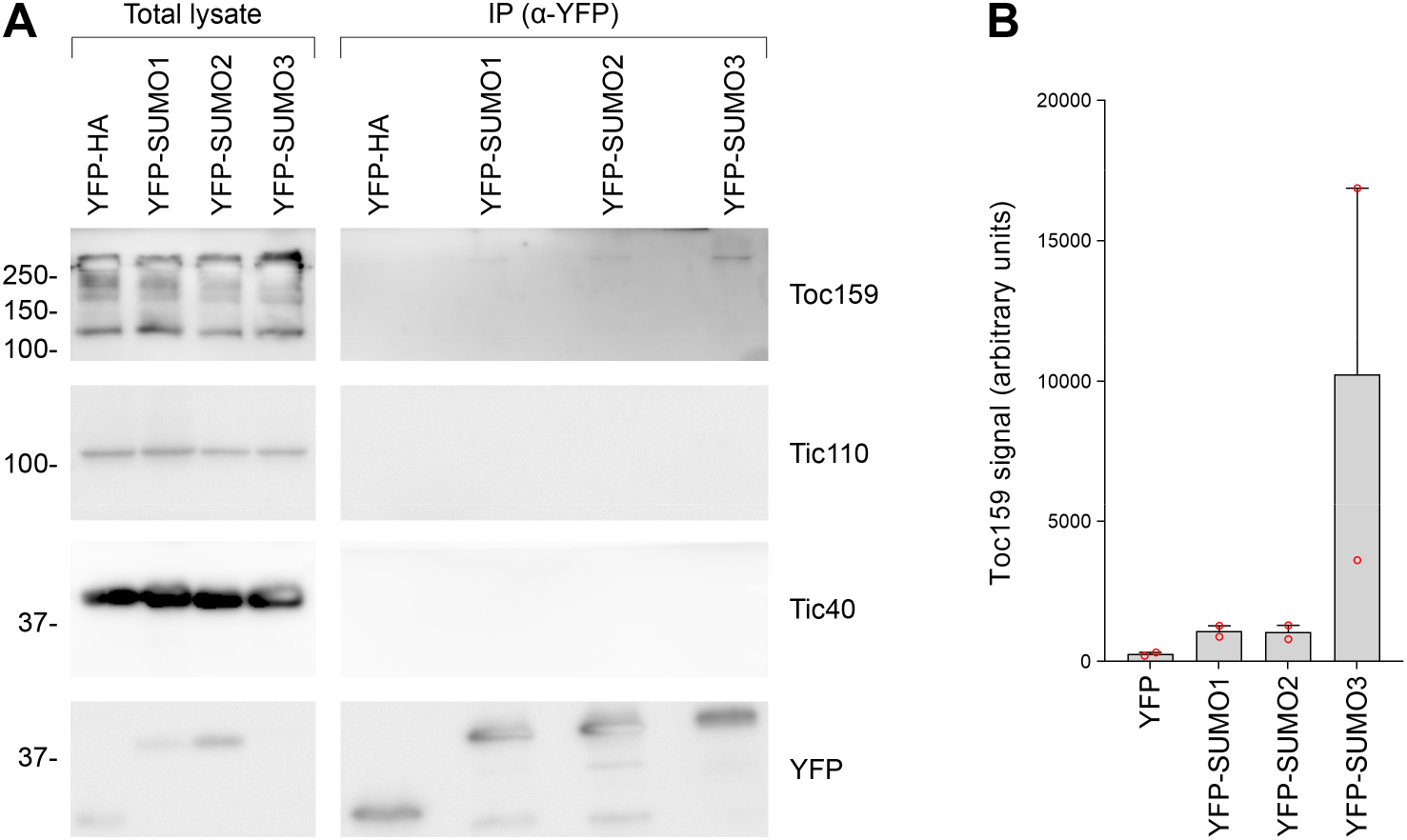
All three SUMO isoforms physically associated with native Toc159, with SUMO3 showing the strongest association. **(A)** The immunoprecipitation experiment shown in Figure 5B was repeated. Tic110 was included as an additional negative control. The samples were run on a 10% acrylamide gel for 1 hour, as opposed to an 8% acrylamide gel for four hours (as in Figure 5B). Migration positions of standards are displayed to the left of the gel images, and sizes are indicated in kDa. The unprocessed membrane images are displayed in Source data 1. **(B)** The Toc159 bands in the eluted fractions in the two experiments, in (A) and in Figure 5B, were quantified. Error bars indicate standard deviation from the mean, and open red circles indicate individual data points.

**Figure 5 – Supplement 4.**
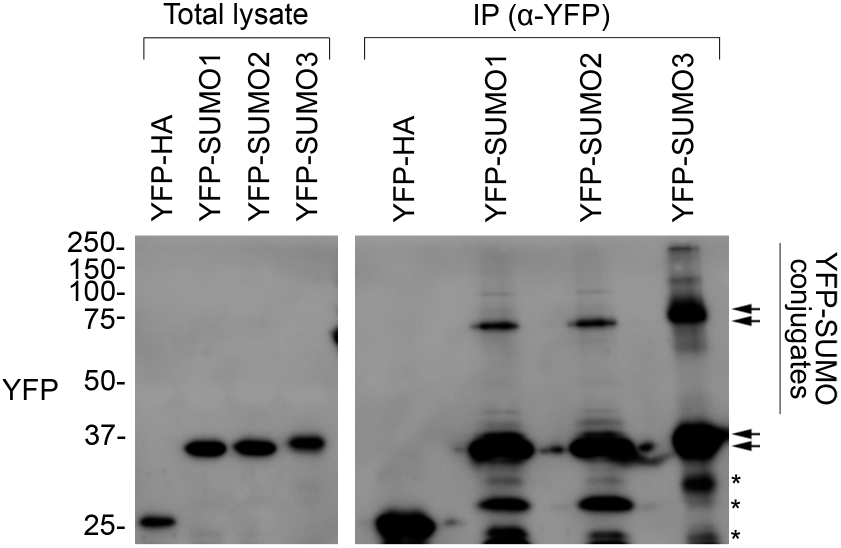
All three YFP-SUMO probes are conjugation-competent. The membrane shown is the same as the one presented in Figure 5B (YFP panel). In this case, the membrane was visualised using a long exposure to aid the detection of weakly abundant protein bands. As well as the YFP-SUMO monomers, additional high molecular bands and smears were seen in the IP samples, indicating the capture of large numbers of SUMOylated proteins. This shows that all three YFP-SUMO probes could be successfully conjugated, as expected (Ayaydin and Dasso, 2004). The positions of the YFP-SUMO monomers (lower pair of arrows), and of the various SUMO adducts (vertical bar) potentially including di-SUMO (upper pair of arrows), are indicated. Note that YFP-SUMO3 migrated more slowly than the other two fusions, as expected. The asterisks indicate non-specific bands. Migration positions of standards are displayed to the left of the gel images, and sizes are indicated in kDa.

## Additional files

### Source data 1

### Immunoblot source data

Uncropped immunoblot images for Figures 1 and 5, and their supplements.

### Transparent reporting form

## References and notes

Ahner, A., Gong, X., and Frizzell, R.A. (2013). Cystic fibrosis transmembrane conductance regulator degradation: cross-talk between the ubiquitylation and SUMOylation pathways. FEBS Journal 280, 4430–4438.

Alonso, J.M., Stepanova, A.N., Leisse, T.J., Kim, C.J., Chen, H., Shinn, P., Stevenson, D.K., Zimmerman, J., Barajas, P., Cheuk, R., Gadrinab, C., Heller, C., Jeske, A., Koesema, E., Meyers, C.C., Parker, H., Prednis, L., Ansari, Y., Choy, N., Deen, H., Geralt, M., Hazari, N., Hom, E., Karnes, M., Mulholland, C., Ndubaku, R., Schmidt, I., Guzman, P., Aguilar-Henonin, L., Schmid, M., Weigel, D., Carter, D.E., Marchand, T., Risseeuw, E., Brogden, D., Zeko, A., Crosby, W.L., Berry, C.C., and Ecker, J.R. (2003). Genome-wide insertional mutagenesis of *Arabidopsis thaliana*. Science 301, 653–657.

Aronsson, H., Combe, J., Patel, R., Agne, B., Martin, M., Kessler, F., Jarvis, P. (2010). Nucleotide binding and dimerization at the chloroplast pre-protein import receptor, atToc33, are not essential in vivo but do increase import efficiency. Plant Journal 63, 297–311.

Ayaydin, F., and Dasso, M. (2004). Distinct in vivo dynamics of vertebrate SUMO paralogues. Molecular Biology of the Cell 15, 5208–5218.

Baek, G.H., Cheng, H., Choe, V., Bao, X., Shao, J., Luo, S., and Rao, H. (2013). Cdc48: a swiss army knife of cell biology. Journal of Amino Acids 2013, 183421.

Bauer, J., Chen, K., Hiltbunner, A., Wehrli, E., Eugster, M., Schnell, D., and Kessler, F. (2000). The major protein import receptor of plastids is essential for chloroplast biogenesis. Nature 403, 203–207.

Beauclair, G., Bridier-Nahmias, A., Zagury, J.F., Saib, A., and Zamborlini, A. (2015). JASSA: a comprehensive tool for prediction of SUMOylation sites and SIMs. Bioinformatics 31, 3483–3491.

Berardini, T.Z., Reiser, L., Li, D., Mezheritsky, Y., Muller, R., Strait, E., and Huala, E. (2015). The Arabidopsis information resource: Making and mining the “gold standard” annotated reference plant genome. Genesis 53, 474–485.

Bergink, S., Ammon, T., Kern, M., Schermelleh, L., Leonhardt, H., and Jentsch, S. (2013). Role of Cdc48/p97 as a SUMO-targeted segregase curbing Rad51-Rad52 interaction. Nature Cell Biology 15, 526–532.

Chang, C.C., Tung, C.H., Chen, C.W., Tu, C.H., and Chu, Y.W. (2018). SUMOgo: Prediction of sumoylation sites on lysines by motif screening models and the effects of various post-translational modifications. Scientific Reports 8, 15512.

Citovsky, V., Lee, L.Y., Vyas, S., Glick, E., Chen, M.H., Vainstein, A., Gafni, Y., Gelvin, S.B., and Tzfira, T. (2006). Subcellular localization of interacting proteins by bimolecular fluorescence complementation *in planta*. Journal of Molecular Biology 362, 1120–1131.

Clough, S.J., and Bent, A.F. (1998). Floral dip: a simplified method for *Agrobacterium*-mediated transformation of *Arabidopsis thaliana*. Plant Journal 16, 735–743.

Desterro, J.M., Rodriguez, M.S., and Hay, R.T. (1998). SUMO-1 modification of IkappaBalpha inhibits NF-kappaB activation. Molecular Cell 2, 233–239.

Elrouby, N., and Coupland, G. (2010). Proteome-wide screens for small ubiquitin-like modifier (SUMO) substrates identify Arabidopsis proteins implicated in diverse biological processes. Proceedings of the National Academy of Sciences of the United States of America 107, 17415–17420.

Flores-Pérez, Ú., and Jarvis, P. (2017). Isolation and suborganellar fractionation of Arabidopsis chloroplasts. Methods in Molecular Biology 1511, 45–60.

Grimmer, J., Helm, S., Dobritzsch, D., Hause, G., Shema, G., Zahedi, R.P., and Baginsky, S. (2020). Mild proteasomal stress improves photosynthetic performance in Arabidopsis chloroplasts. Nature Communications 11, 1662.

Hilgarth, R.S., and Sarge, K.D. (2005). Detection of sumoylated proteins. Methods in Molecular Biology 301, 329–338.

Hirsch, S., Muckel, E., Heemeyer, F., von Heijne, G., and Soll, J. (1994). A receptor component of the chloroplast protein translocation machinery. Science 266, 1989–1992.

Huang, W.H., Ling, Q.H., Bedard, J., Lilley, K., and Jarvis, P. (2011). In vivo analyses of the roles of essential Omp85-related proteins in the chloroplast outer envelope membrane. Plant Physiology 157, 147–159.

Inaba, T., Alvarez-Huerta, M., Li, M., Bauer, J., Ewers, C., Kessler, F., and Schnell, D.J. (2005). Arabidopsis Tic110 is essential for the assembly and function of the protein import machinery of plastids. Plant Cell 17, 1482–1496.

Ishida, T., Fujiwara, S., Miura, K., Stacey, N., Yoshimura, M., Schneider, K., Adachi, S., Minamisawa, K., Umeda, M., and Sugimoto, K. (2009). SUMO E3 ligase HIGH PLOIDY2 regulates endocycle onset and meristem maintenance in Arabidopsis. Plant Cell 21, 2284–2297.

Jarvis, P. (2008). Targeting of nucleus-encoded proteins to chloroplasts in plants. New Phytologist 179, 257–285.

Jarvis, P., and Lopez-Juez, E. (2013). Biogenesis and homeostasis of chloroplasts and other plastids. Nature Reviews Molecular Cell Biology 14, 787–802.

Jarvis, P., Chen, L.J., Li, H.M., Pete, C.A., Fankhauser, C., and Chory, J. (1998). An *Arabidopsis* mutant defective in the plastid general protein import apparatus. Science 282, 100–103.

Karimi, M., Inze, D., and Depicker, A. (2002). GATEWAY vectors for *Agrobacterium*-mediated plant transformation. Trends in Plant Science 7, 193–195.

Karimi, M., De Meyer, B., and Hilson, P. (2005). Modular cloning in plant cells. Trends Plant Science 10, 103–105.

Kasmati, A.R., Töpel, M., Patel, R., Murtaza, G., and Jarvis, P. (2011). Molecular and genetic analyses of Tic20 homologues in *Arabidopsis thaliana* chloroplasts. Plant Journal 66, 877–889.

Kessler, F., Blobel, G., Patel, H.A., and Schnell, D.J. (1994). Identification of two GTP-binding proteins in the chloroplast protein import machinery. Science 266, 1035–1039.

Kovacheva, S., Bedard, J., Patel, R., Dudley, P., Twell, D., Rios, G., Koncz, C., and Jarvis, P. (2005). *In vivo* studies on the roles of Tic110, Tic40 and Hsp93 during chloroplast protein import. Plant Journal 41, 412–428.

Kurepa, J., Walker, J.M., Smalle, J., Gosink, M.M., Davis, S.J., Durham, T.L., Sung, D.Y., and Vierstra, R.D. (2003). The small ubiquitin-like modifier (SUMO) protein modification system in Arabidopsis. Accumulation of SUMO1 and −2 conjugates is increased by stress. Journal of Biological Chemistry 278, 6862–6872.

Lee, S., Lee, D.W., Lee, Y., Mayer, U., Stierhof, Y.D., Jurgens, G., and Hwang, I. (2009). Heat shock protein cognate 70-4 and an E3 ubiquitin ligase, CHIP, mediate plastid-destined precursor degradation through the ubiquitin-26S proteasome system in *Arabidopsis*. Plant Cell 21, 3984–4001.

Lee, Y.J., Kim, D.H., Kim, Y.W., and Hwang, I. (2001). Identification of a signal that distinguishes between the chloroplast outer envelope membrane and the endomembrane system in vivo. Plant Cell 13, 2175–2190.

Liebelt, F., and Vertegaal, A.C. (2016). Ubiquitin-dependent and independent roles of SUMO in proteostasis. American Journal of Physiology Cell Physiology 311, C284–C296.

Ling, Q., and Jarvis, P. (2015). Regulation of chloroplast protein import by the ubiquitin E3 ligase SP1 is important for stress tolerance in plants. Current Biology 25, 2527–2534.

Ling, Q., Huang, W., and Jarvis, P. (2011). Use of a SPAD-502 meter to measure leaf chlorophyll concentration in Arabidopsis thaliana. Photosynthesis Research 107, 209–214.

Ling, Q., Huang, W., Baldwin, A., and Jarvis, P. (2012). Chloroplast biogenesis is regulated by direct action of the ubiquitin-proteasome system. Science 338, 655–659.

Ling, Q., Broad, W., Trösch, R., Töpel, M., Demiral Sert, T., Lymperopoulos, P., Baldwin, A., and Jarvis, R.P. (2019). Ubiquitin-dependent chloroplast-associated protein degradation in plants. Science 363.

Liu C., Yu H., Li L. (2019). SUMO modification of LBD30 by SIZ1 regulates secondary cell wall formation in *Arabidopsis thaliana*. PLOS Genetics 15, e1007928.

Miura, K., Rus, A., Sharkhuu, A., Yokoi, S., Karthikeyan, A.S., Raghothama, K.G., Back, D., Koo, Y.D., Jin, J.B., Bressan, R.A., Yun, D.J., and Hasegawa, P.M. (2005). The Arabidopsis SUMO E3 ligase SIZ1 controls phosphate deficiency responses. Proceedings of the National Academy of Sciences of the United States of America 102, 9734–9734.

Nie, M., Aslanian, A., Prudden, J., Heideker, J., Vashisht, A.A., Wohlschlegel, J.A., Yates, J.R., and Boddy, M.N. (2012). Dual recruitment of Cdc48 (p97)-Ufd1-Npl4 ubiquitin-selective segregase by small ubiquitin-like modifier protein (SUMO) and ubiquitin in SUMO-targeted ubiquitin ligase-mediated genome stability functions. Journal of Biological Chemistry 287, 29610–29619.

Perry, S.E., and Keegstra, K. (1994). Envelope membrane proteins that interact with chloroplastic precursor proteins. Plant Cell 6, 93–105.

Saracco, S.A., Miller, M.J., Kurepa, J., and Vierstra, R.D. (2007). Genetic analysis of SUMOylation in Arabidopsis: conjugation of SUMO1 and SUMO2 to nuclear proteins is essential. Plant Physiology 145, 119–134.

Schneider, C.A., Rasband, W.S., and Eliceiri, K.W. (2012). NIH Image to ImageJ: 25 years of image analysis. Nature Methods 9, 671–675.

Schnell, D.J., Kessler, F., and Blobel, G. (1994). Isolation of components of the chloroplast protein import machinery. Science 266, 1007–1012.

Shanmugabalaji, V., Chahtane, H., Accossato, S., Rahire, M., Gouzerh, G., Lopez-Molina, L., and Kessler, F. (2018). Chloroplast biogenesis controlled by DELLA-TOC159 interaction in early plant development. Current Biology 28, 2616–2623 e2615.

Sowden, R.G., Watson, S.J., and Jarvis, P. (2018). The role of chloroplasts in plant pathology. Essays in Biochemistry 62, 21–39.

Sriramachandran, A.M., and Dohmen, R.J. (2014). SUMO-targeted ubiquitin ligases. Biochimica et Biophysica Acta 1843, 75–85.

Tranel, P.J., Froehlich, J., Goyal, A., and Keegstra, K. (1995). A component of the chloroplastic protein import apparatus is targeted to the outer envelope membrane via a novel pathway. The EMBO Journal 14, 2436–2446.

Tzfira, T., Tian, G.W., Lacroix, B., Vyas, S., Li, J., Leitner-Dagan, Y., Krichevsky, A., Taylor, T., Vainstein, A., and Citovsky, V. (2005). pSAT vectors: a modular series of plasmids for autofluorescent protein tagging and expression of multiple genes in plants. Plant Molecular Biology 57, 503.

van den Burg, H.A., Kini, R.K., Schuurink, R.C., and Takken, F.L.W. (2010). *Arabidopsis* small ubiquitin-like modifier paralogs have distinct functions in development and defense. Plant Cell 22, 1998–2016.

Watson, S.J., Sowden, R.G., and Jarvis, P. (2018). Abiotic stress-induced chloroplast proteome remodelling: a mechanistic overview. Journal of Experimental Botany 69, 2773–2781.

Wu, F.H., Shen, S.C., Lee, L.Y., Lee, S.H., Chan, M.T., and Lin, C.S. (2009). Tape-Arabidopsis Sandwich - a simpler Arabidopsis protoplast isolation method. Plant Methods 5, 16.

Zhao, Q., Xie, Y., Zheng, Y., Jiang, S., Liu, W., Mu, W., Liu, Z., Zhao, Y., Xue, Y., and Ren, J. (2014). GPS-SUMO: a tool for the prediction of sumoylation sites and SUMO-interaction motifs. Nucleic Acids Research 42, W325–W330.

