## Supplementary material for "Crosstalk between chloroplast protein import and the SUMO system revealed through genetic and molecular investigation": Source data 1

Immunoblot source data for:

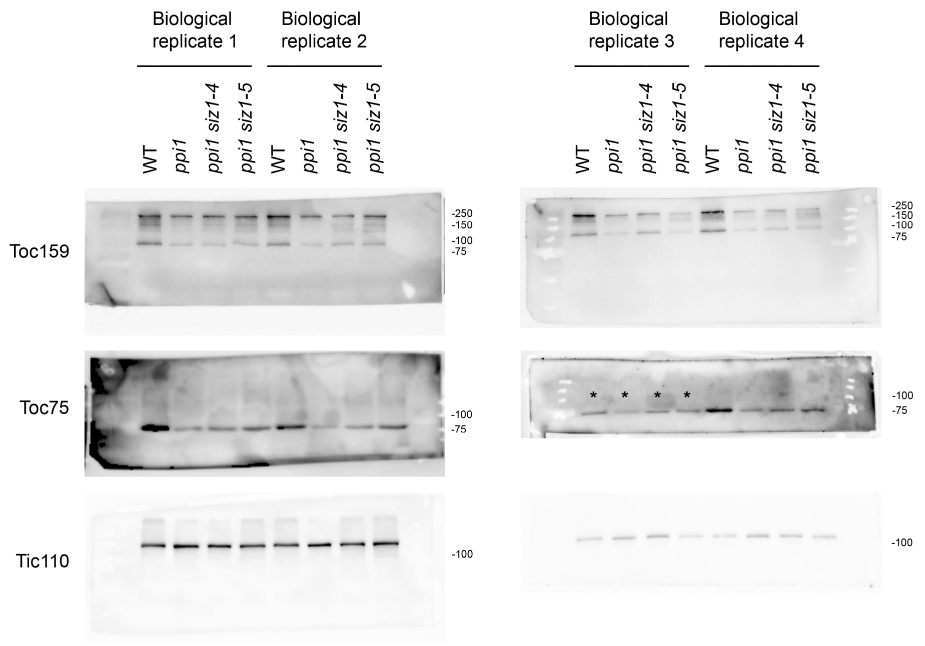


**Source data for Figures 1I, 1J and 1K.**

Protein samples were taken from the plants pictured in Figure 1 – Supplement 1C on the day of photography. Four independent protein samples were taken from each genotype (four biological replicates). The asterisks indicate that the samples were not included in the Figure 1K quantification; these samples were excluded because the WT Toc75 band was unusually weak, possibly due to a transfer problem in this part of the membrane.

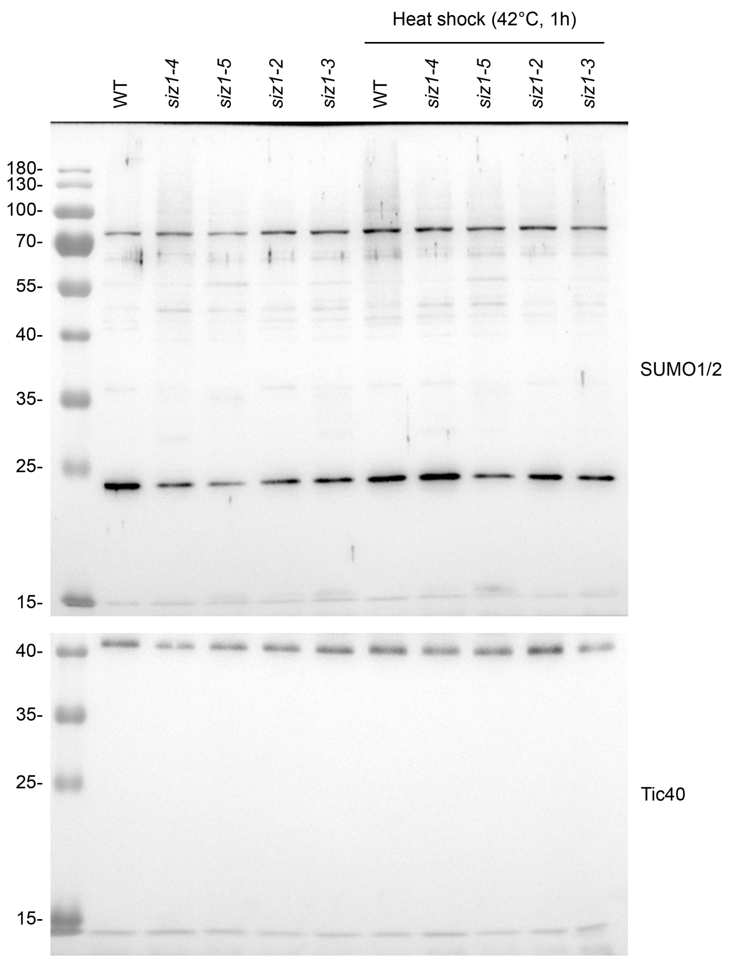


**Source data for Figure 1 – Supplement 3.**

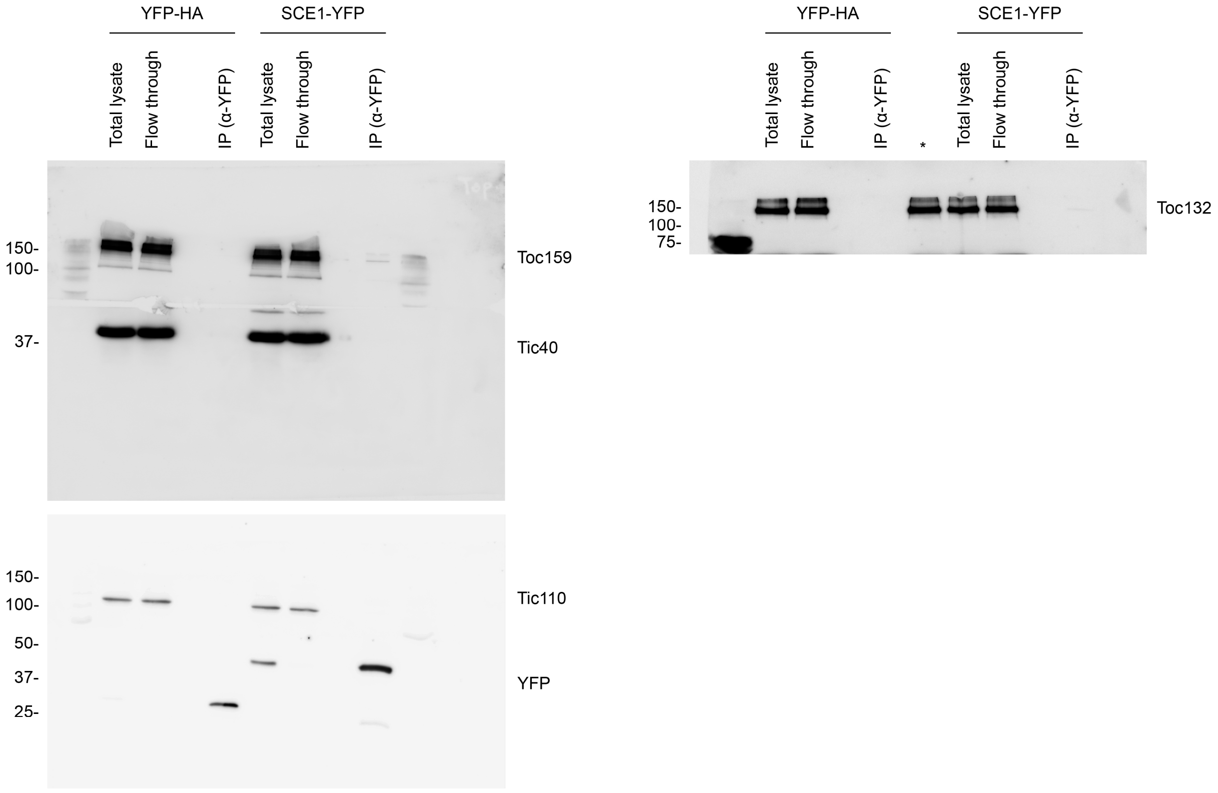


**Source data for Figure 5A.**

The asterisk indicates that the lane was omitted from the processed figure. This lane contained the ‘total lysate’ sample, which was loaded twice in error.

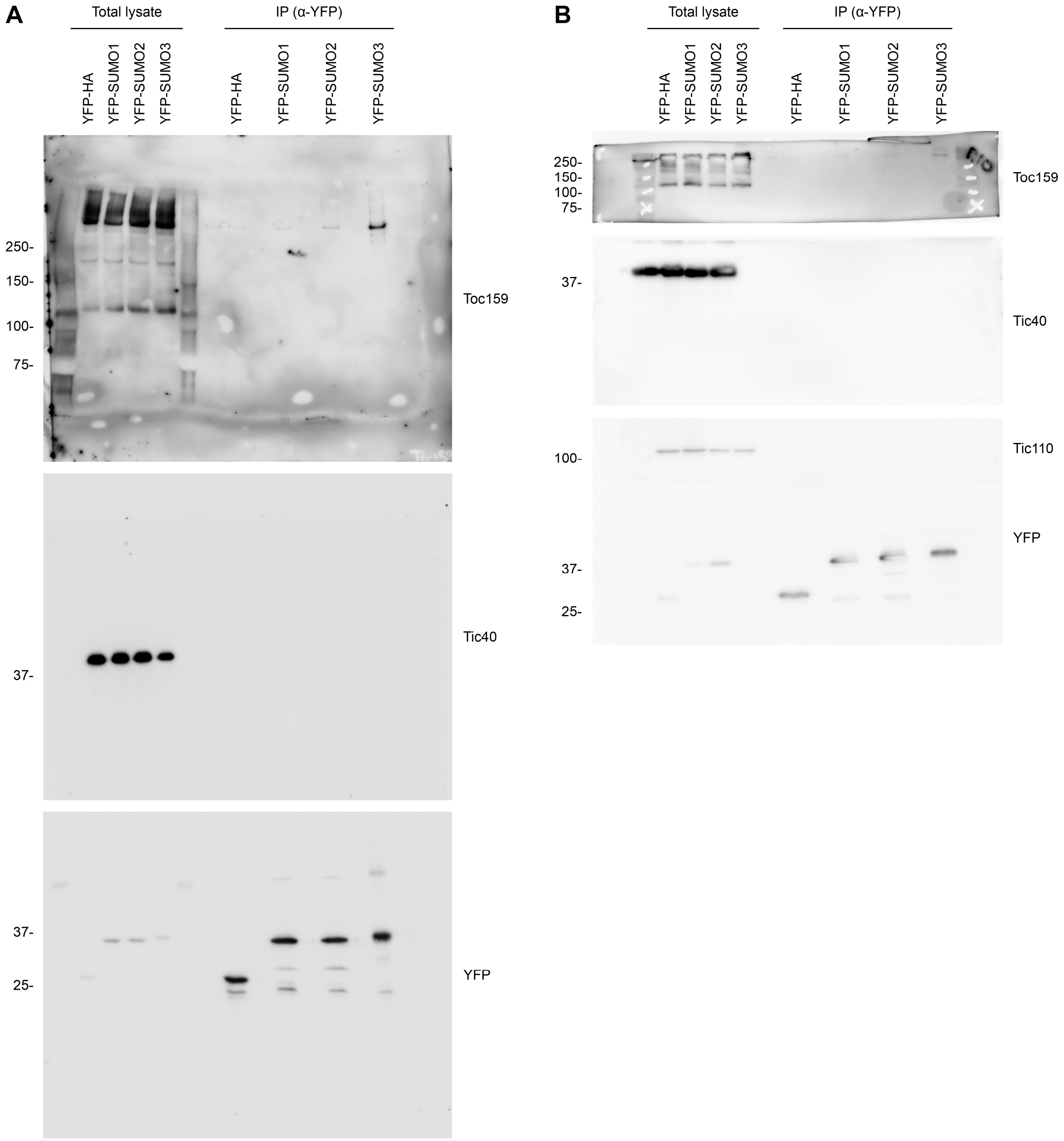


**Source data for Figure 5B and Figure 5 – Supplement 3.**

The data in **(A)** support Figure 5B, and the data in **(B)** support Figure 5 – Supplement 3.

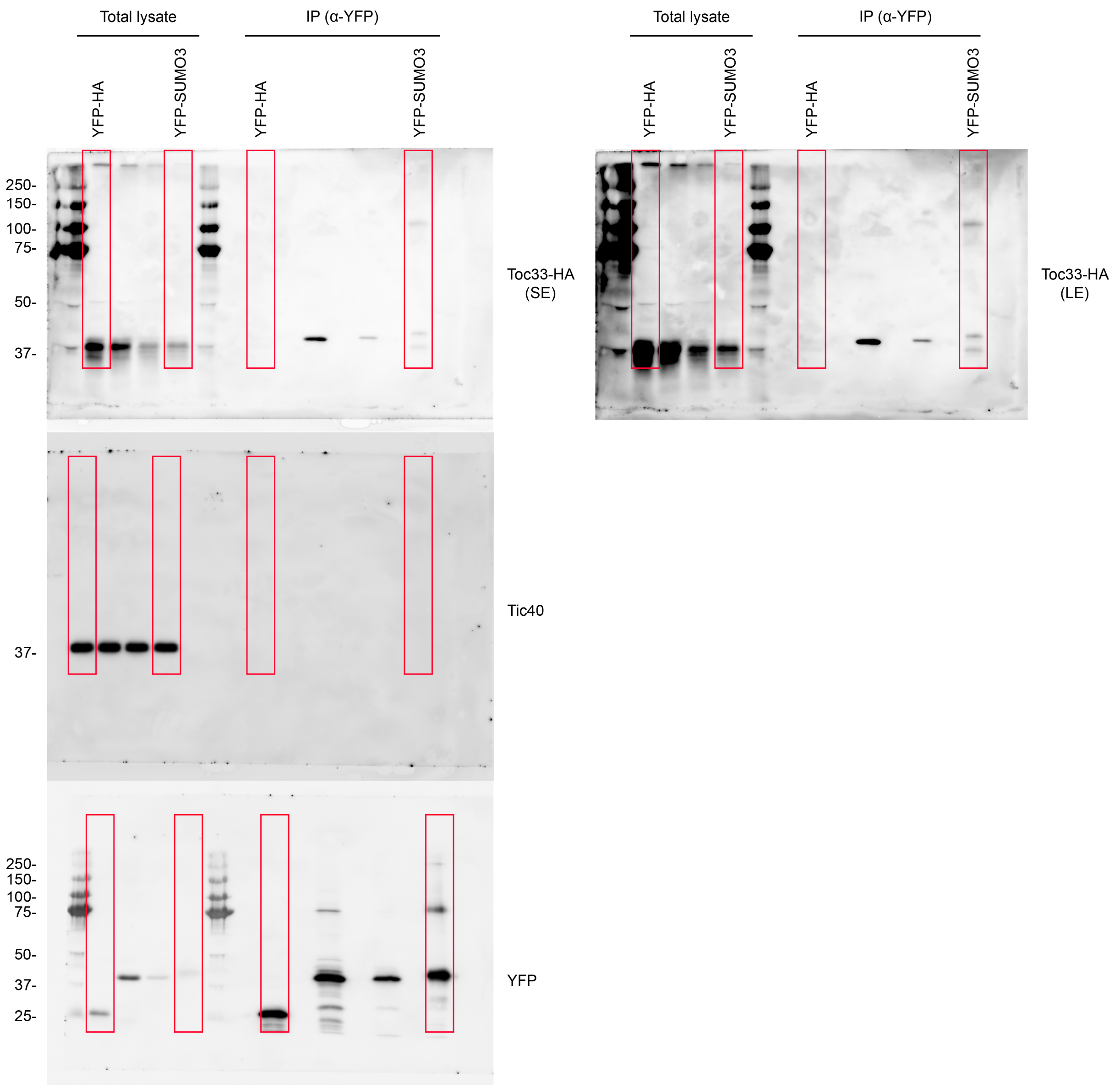


**Source data for Figure 5C.**

The red boxes indicate the relevant lanes.
